# Inter-individual variation in human cortical cell type abundance and expression

**DOI:** 10.1101/2022.10.07.511366

**Authors:** Nelson Johansen, Saroja Somasundaram, Kyle J. Travaglini, Anna Marie Yanny, Maya Shumyatcher, Tamara Casper, Charles Cobbs, Nick Dee, Richard Ellenbogen, Manuel Ferreira, Jeff Goldy, Junitta Guzman, Ryder Gwinn, Daniel Hirschstein, Nikolas L. Jorstad, C. Dirk Keene, Andrew Ko, Boaz P. Levi, Jeffrey G. Ojemann, Thanh Pham, Nadiya Shapovalova, Daniel Silbergeld, Josef Sulc, Amy Torkelson, Herman Tung, Kimberly Smith, Ed S. Lein, Trygve E. Bakken, Rebecca D. Hodge, Jeremy A. Miller

**Affiliations:** Allen Institute for Brain Science; Seattle, USA; Swedish Neuroscience Institute; Seattle, USA; University of Washington, Neurological Surgery; Seattle, USA; University of Washington, Department of Laboratory Medicine and Pathology; Seattle, USA

## Abstract

Single cell transcriptomic studies have identified a conserved set of neocortical cell types from small post-mortem cohorts. We extend these efforts by assessing cell type variation across 75 adult individuals undergoing epilepsy and tumor surgeries. Nearly all nuclei map to one of 125 robust cell types identified in middle temporal gyrus, but with varied abundances and gene expression signatures across donors, particularly in deep layer glutamatergic neurons. A minority of variance is explainable by known factors including donor identity and small contributions from age, sex, ancestry, and disease state. Genomic variation was significantly associated with variable expression of 150-250 genes for most cell types. Thus, human individuals display a highly consistent cellular makeup, but with significant variation reflecting donor characteristics, disease condition, and genetic regulation.

**One-Sentence Summary:** Inter-individual variation in human cortex is greatest for deep layer excitatory neurons and largely unexplainable by known factors.

## Main Text

The human brain displays well-established inter-individual variability in regional activity, morphology, and connectivity (*1*), with neurotypical brains varying up to two-fold in size and showing a wide range of associated changes in brain shape (*2*). This variation necessitates the use of common coordinate frameworks where individual human brain maps are morphed to fit a representative average for direct comparisons across individuals (*3*). Such methods have shown that brain structure in humans relates to many demographic and behavior variables across a large cohort of young people (*4*), and that genetic variants impact the structure of subcortical areas (*5*). Furthermore, functional networks defined with resting-state functional magnetic resonance imaging (fMRI) associate with ion channel and synaptic gene networks in neurotypical adult donors (*6*), providing a link between functional and transcriptomic networks. Resting-state fMRI can also be tied to population variation of the underlying genome, where common genetic variants have been shown to influence intrinsic brain activity in a manner correlated with brain disorders such as schizophrenia and major depressive disorder (*7*), further extending the causal chain.

Despite this extensive body of imaging data demonstrating inter-individual variability in human brains, we lack an understanding of population differences in cell type distributions and gene expression and how these differences are impacted by genetic, environmental, and demographic factors. Neuronal proportions are established during prenatal development and stable through adulthood and may contribute to behavioral variation, whereas non-neuronal proportions can change in response to the environment, such as microglial proliferation in response to inflammatory stress (*8*). Cellular gene expression profiles reflect the developmental origins, life history, and current activity of a cell. A previous study demonstrated stable gene expression patterning across brain regions in six human donors using microarray profiling of bulk tissue dissections (*9*), but the consistency or variability of such gene networks across cell types and larger human cohorts is unknown (*10*, *11*).

Single nucleus transcriptomics (snRNA-seq) provides a major advance over previous genomic assays by separating gene expression changes due to differences in expression levels from changes due to differences in cell type abundances across individuals (*12*–*15*). Previous snRNA-seq studies provide hints about anticipated variation in gene expression and cell type abundance. Studies of small cohorts (N=3-8) of young adult human brain donors show strong inter-individual concordance, as cells from different donors co-cluster by cell type without the need for data integration methods (*16*, *17*). In contrast, high variation in cell type proportions and gene expression are seen in larger studies of Alzheimer’s disease and other neurodegenerative diseases, which can largely be explained by disease phenotypes (*18*–*20*). These results complement earlier microarray studies that demonstrate increased transcriptional heterogeneity in human brain with age and associated with many gene pathways (*21*). Furthermore, gene expression variation differs by cell type. In mouse, glutamatergic types show greater gene expression changes across cortical regions than GABAergic types, with unbiased clustering identifying distinct glutamatergic types in primary visual and anterior lateral motor cortices (*22*). Similarly, supragranular glutamatergic neurons are highly variable across several axes, including cell type, cortical depth, donor, and species (*16*, *17*, *23*). However, the extent of such variation across many individuals is not clear.

Heterogeneity in gene expression across individuals can arise as a function of biological factors such as age, sex, or ancestry, as well as the presence of genetic variants that modulate expression. Combined genotype and snRNA-seq data in even small cohorts permit investigation of the functional effects of disease-associated variants and identification of genes whose expression is under genetic control through expression quantitative trait locus (eQTL) analysis. Existing studies that perform eQTL analysis using bulk RNA-seq or snRNA-seq provide insights into brain-specific eQTLs (*24*, *25*), some of which are associated with diseases like Alzheimer’s disease (*20*, *26*, *27*). However, the current resolution of cell type-specific eQTL analysis is limited to the broadest cell types and the extent to which genetic variants can modulate expression within finer cell type annotation remains unexplored.

In short, additional work is needed to determine the degree of inter-individual variability in transcriptional programs in human brain and to assess how this variability relates to cell types and genotypic and phenotypic variables. Here we investigated variation in cortical cellular abundance and gene expression across 75 adult individuals, providing a detailed view at how cell type variation reflects donor characteristics, disease condition, and the underlying genomic landscape.

## Results

### High quality tissue from neurosurgical resections

We collected overlying cortical tissue from 75 adult individuals undergoing neurosurgery for intractable epilepsy or removal of tumors (**Table S1**). Most tissue was from middle temporal gyrus (MTG) removed to gain access to underlying hippocampal tissues during epilepsy surgery, and all but four of the remaining tissue samples were derived from frontal cortex (FRO) (**Fig. 1a**). Approximately two thirds of donors underwent epilepsy surgery, and these individuals were generally younger than those with tumors. Males represented just over half of the donors from each medical condition.

**Fig. 1.**
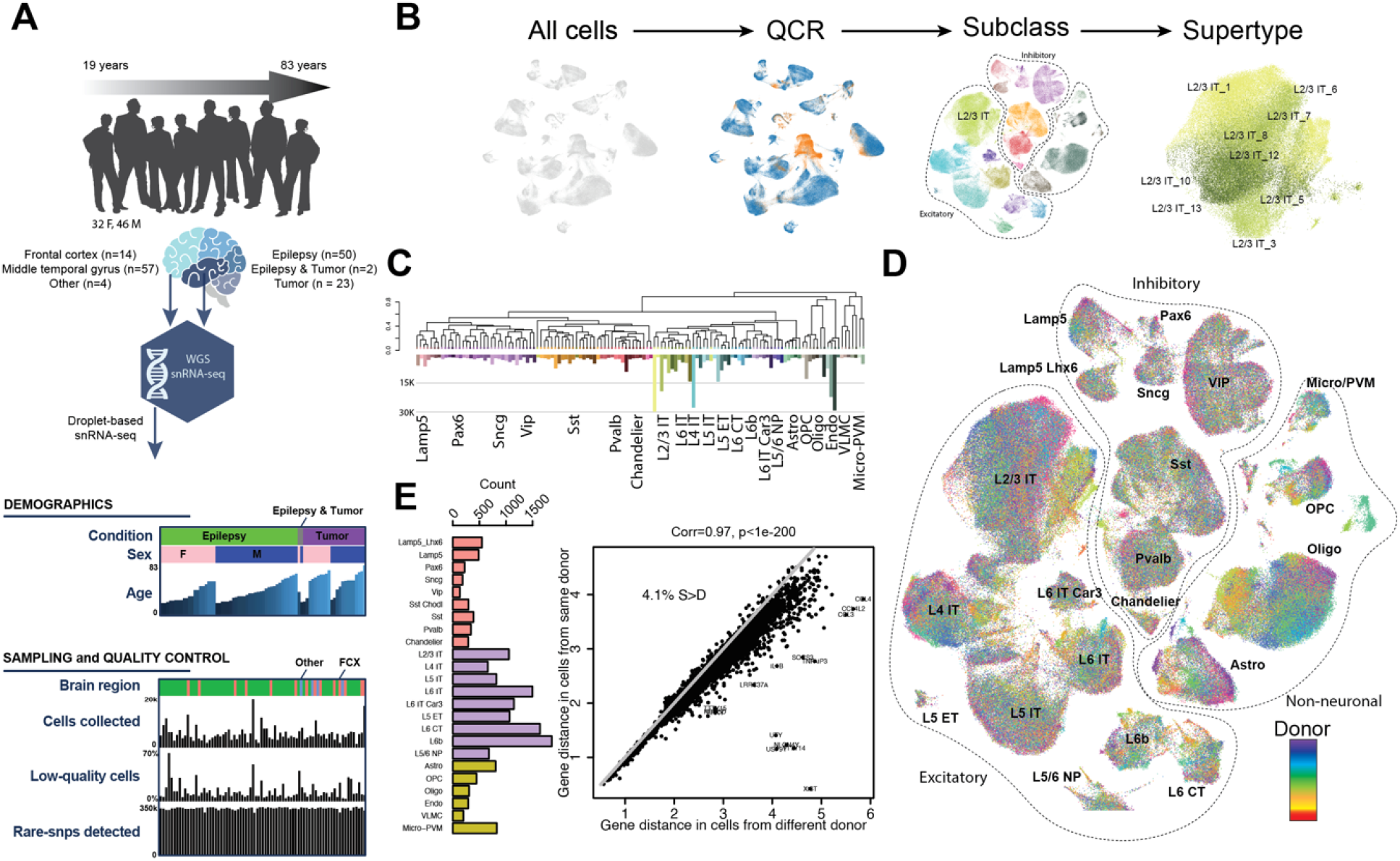
Study design and overall gene expression variation in neurosurgical cohort. (**A**) Summary of data types, individuals, demographics, and quality metrics in this study. (**B**) Schematic of quality control and cell type assignment for single nuclei collected in this study. (**C**) (top) Dendrogram of reference supertypes from this study (identical to the taxonomy from sea-ad.org). (bottom) Number of cells from this study mapped to each supertype. Most GABAergic types are rare, while some superficial IT and oligodendrocyte types are quite common. (**D**) Low dimensional (UMAP) representation of all cells from this study, color-coded by donor and labeled by subclass. Dotted lines separate broad cell classes, which are also labeled. The same UMAP representation is used in Fig 1B, 1D, and 2C. (**E**) (left) Number of genes with significantly higher distance between cells from different donors as compared to cells from the same donor (x-axis) for each subclass (y-axis). FDR<0.00494 was chosen such that no genes have significantly higher intra-donor cell distance. (right) Scatterplot showing median inter-donor (x-axis) vs. intra-donor (y-axis) distance for each gene expressed in at least one type (points). The top 15 most variable genes are shown, which include several sex chromosome and immune-response genes.

**Fig. 2.**
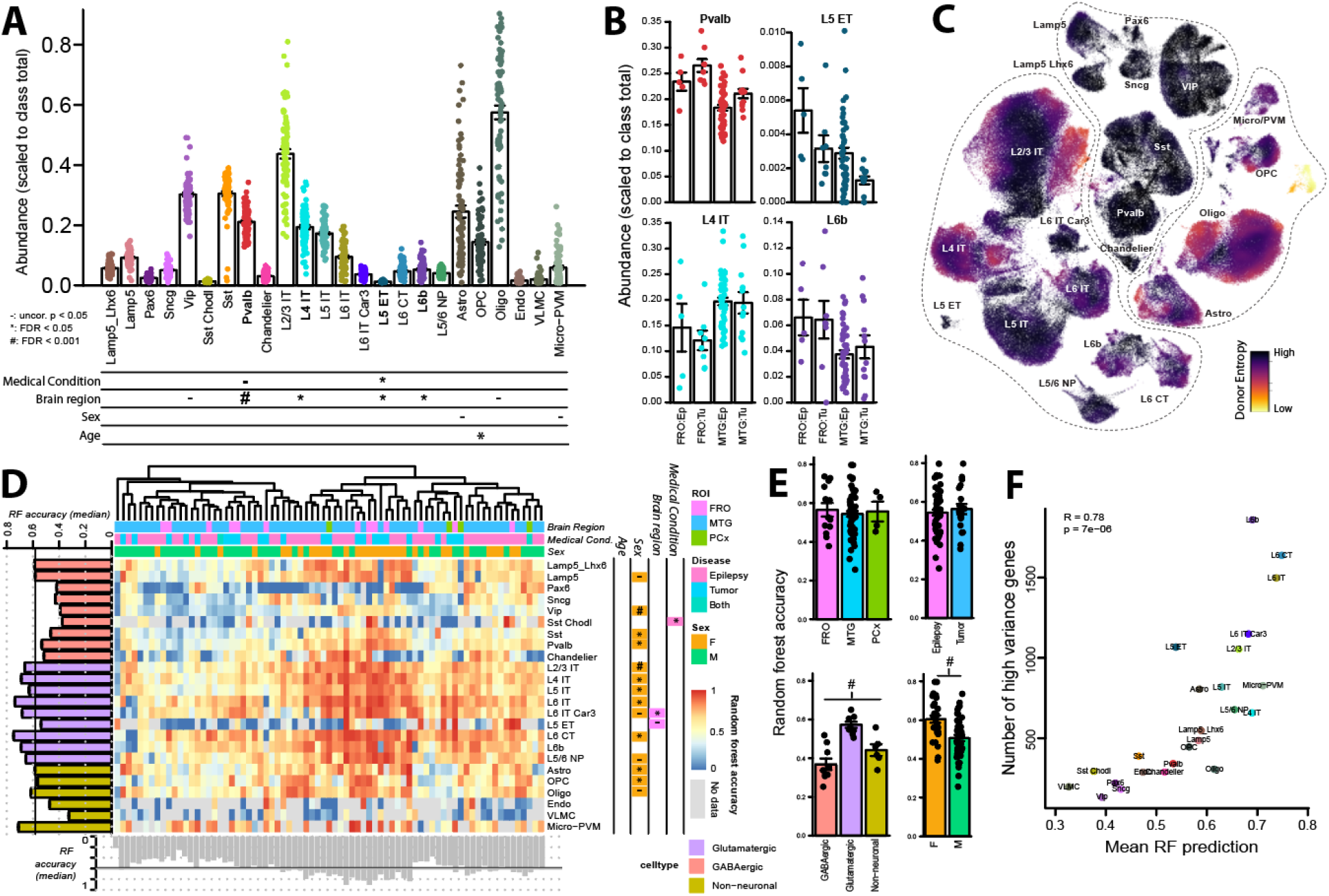
Cell type-specific differences in abundance and random forest predictions across donors. (**A**) (top) Fraction of cells from each subclass collected from each donor, scaled to the total number of cells from each class (GABAergic neuron, glutamatergic neuron, non-neuron). For bar plots in all panels, points represent values from individual donors, with bars showing mean +/− standard error. (bottom) Significant associations between subclass abundances and the listed metadata when performing a linear model with covariates for medical condition, brain region, sex, age, and batch (−,*,# as indicated). (**B**) Bar plots showing abundances after dividing donors by medical condition and brain region for select subclasses with significant changes in at least one of these metrics. (**C**) Lower donor entropy in GABAergic interneurons than other types. UMAP as in Figure 1, color-coded to show donor entropy, which is a measure of the number of donors represented in nearby nuclei. (**D**) Random forest (RF) predictability of cells differs by cell type and donor characteristics. Heatmap shows fraction of nuclei of a given subclass (x-axis) correctly assigned to a given donor (y-axis) using random forest classification with 75% training / 25% test strategy. Bar plots on the x- and y- axes show the median fraction of cells correctly classified by donor and subtype, respectively. Cell types and donors colored as indicated in the legend. Significant associations between classification accuracies and the listed metadata when performing the same linear model as in A are shown, additionally color-coded by direction of maximum accuracy. (**E**) Bar plots showing median random forest accuracies after dividing donors or cell types by listed metrics identify higher prediction accuracy in glutamatergic than other subclasses, and in females as compared with males. (**F**) High correlation between the number of high variance genes per subclass (y-axis, **Fig. 1E**) and the mean RF predication accuracy.

Tissue was collected for droplet-based snRNA-seq and whole genome sequencing (WGS) from adjacent sections to those processed for patch-seq studies (*23*) following a published protocol (*28*) (**Methods**). We previously showed that resected tissue is largely neurotypical, with no obvious relationship between electrophysiology or morphology and pathology, sex, or age (*23*), or time in patch clamp solution (*28*). Histological assessments of neurodegeneration, gliosis, and tumor infiltration show minimal indicators of pathology (*23*). Finally, gene expression from neurosurgical and postmortem tissue has high fidelity, as nuclei from surgical tissues show relatively few gene expression changes as compared to postmortem tissues (*17*), and patch-seq cells can be mapped to reference transcriptomic classifications built using predominantly postmortem nuclei with high confidence (*23*).

### Nuclei from all donors mapped to robust MTG reference taxonomy

To assess cell type variation across individuals, it is critical to have a high-confidence reference taxonomy. Here we use a human MTG taxonomy created from ~150,000 nuclei collected from postmortem tissue of five neurotypical donors as part of the Seattle Alzheimer’s Disease Brain Cell Atlas (SEA-AD; sea-ad.org). These cell types are also used in SEA-AD and mapping to the same reference will allow future comparisons between cells from younger and aged donors. 125 “supertypes” were defined as a set of fine-grained cell type annotations for single nucleus expression data that could be reliably predicted with an F1 score greater than 0.7 on held-out reference data (*29*) (**Methods**).

Nuclei from each donor underwent basic filtering on the number of genes detected (>500) and doublet scores (<0.3) and were then assigned to cell types iteratively by first mapping to cell “class” (e.g., glutamatergic neuron), followed by “subclass” (e.g., layer 2/3 intratelencephalic glutamatergic neuron [L2/3 IT]), and finally by supertype (e.g., L2/3 IT_6; **Fig. 1b**; **Fig. S1**, **Methods**). Next, to define high quality nuclei, we developed a novel algorithm for automated quality control that can be applied to snRNA-seq data from one or more libraries (QCR; **Methods**) and used this to flag nuclei for potential filtering. These flags were then used as a baseline for manual filtering of low-quality cells and doublets in an instance of CELLxGENE (*30*) that included these data and nuclei from the SEA-AD reference. For most individuals <15% of nuclei were excluded, with >50% of nuclei filtered in three cases, resulting in an average of 4,597 (IQR: 3,344-5,396) nuclei collected per individual after controlling for RNA quality (**Fig. 1a**).

To address the relative sparsity of NeuN-nuclei in this reference and the known heterogeneity of non-neuronal cell types, we extended the set of non-neuronal supertypes by reclustering these nuclei alongside non-neuronal nuclei from the 75 individuals in this study, which resulted in an additional set of 6 Astrocyte, 1 Micro-PVM, 3 Oligodendrocyte, and 1 OPC transcriptionally distinct populations (**Fig. S1**). As with postmortem donors, most cell types were rare except for several neuronal (IT) and glial types (**Fig. 1c**). Due to the overall sparsity of supertypes as well as potential gene expression differences in glutamatergic types across brain regions (*22*, *31*) we focus many of our comparisons on high fidelity subclass assignments.

### Glutamatergic neurons show highest inter-donor variability

To gain an initial understanding of cell-to-cell variability we plotted a UMAP of the entire data set using the most variable genes without performing additional data integration and color-coding by different metadata. Cells show nearly perfect separation by subclass, often dividing into distinct islands (**Fig. 1b**), indicating that cell types are highly consistent across donors at this level of resolution. Within each subclass, most supertypes also show visual separation indicating that these higher resolution cell type assignments are relatively robust across donors (**Fig. S1**). In contrast, inter-individual variability is highly dependent on cell type. Inhibitory neurons are well mixed by donors, whereas glutamatergic neurons and some non-neurons show spotting and banding by donor (**Fig. 1d**), indicating that at least some donors have gene expression profiles different from other donors for these cell types. Indeed, we find a small set of non-neuronal types with distinct transcriptional signatures that are comprised of nuclei from only two donors with tumors (**Fig. 1d**). Further examination of available histology showed evidence of infiltration of abnormal cells into donor tissue samples, suggesting that these transcriptionally distinct cells may reflect disease-related cell states.

To examine which genes are most variable across individuals, we divided cells by subclass and compared the gene distance between pairs of nuclei from the same donor against pairs of nuclei collected from different donors to define “high variance genes”. Using an FDR threshold such that no genes showed larger within-than between-donor distance (FDR < 0.00494), all glutamatergic subclasses had more high variance genes than all GABAergic and most non-neuronal types (**Fig. 1e**). Specifically, glutamatergic types in layer 6 had the most high-variance genes (N >= 1149 for L6 IT Car3, L6 IT, L6 CT, L6b), although after scaling by the number of genes expressed, Microglia were the most diverse across individuals (**Fig. S2a**). In contrast, Vip, Sncg, and Pax6 GABAergic subclasses and non-neural types (VLMC and endothelial cells) had the fewest such genes (N<=289). Gene counts were not correlated with the number of cells per subclass (**Fig. S2b**), suggesting that this is a biological rather than a technical result. Many high variance genes have clear biological contexts (**Fig. 1e**; **Fig. S2c-d**). Sex genes, including genes on the Y chromosome (UTY, TTTY14) and XIST, a key regulator of X-chromosome inactivation in females (*32*), are among the most highly variable genes across nearly all subclasses. Similarly variable across most subclasses were immediate early genes (FOS, JUN, JUND), the expression of which can change rapidly in response extracellular stimuli (*33*), and which we previously showed are upregulated in neurosurgical tissue relative to postmortem tissue (*17*). Finally, genes associated with the overall amount of nuclear RNA, including housekeeping genes (GAPDH, ACTB) and MALAT1, the most highly expressed and selective transcript in the nucleus (*34*) that also regulates synaptic density (*35*), show high variance across donors in most subclasses, suggesting that overall transcription may also vary by individual.

Next, we performed gene ontology (GO) enrichment analysis separately for each subclass (**Fig. S2e**). Variable genes in glial types were significantly enriched for “glutamatergic synapse” genes (FDR<=1.1E-6 for oligodendrocytes, OPCs, and astrocytes), which capture known genes involved in neuron-glia and glia-glia signaling and which are also the most variable genes when comparing human to non-human primate (*36*). Inflammatory response genes were overrepresented among highly variable genes selectively in microglia (FDR = 3.9E-14), suggesting an increase in reactive microglia in a subset of individuals. These include marker genes CCL3, CCL4, CCL4L2, IL1B (**Fig. S2d**), which were among the most highly variable in this study (**Fig. 1e**). Histology available for a partially overlapping set of individuals found moderate IBA1 reactivity in a small but significant fraction of individuals (see Extended data Figure 1 from (*23*)), supporting this result. Few GO categories were enriched for neuronal subclasses (**Fig. S2e**), suggesting that the set of variable genes span many biological functions, perhaps not surprisingly given the many potential sources of inter-individual variation (e.g., brain region, sex, age, and medical condition). Together these results indicate significant and cell type-dependent inter-individual variability, whose origins we more carefully dissect below.

### Consistent but more variable metrics in neurosurgical than postmortem tissue

Given differences in medical conditions, individual life experiences, and tissue collection protocols, we directly assessed changes in gene expression and cellular abundance between neurosurgical and postmortem tissue samples. To limit extraneous sources of variation in this analysis, we only considered 45 individuals undergoing epilepsy surgery in MTG and compared them to the three donors from the reference MTG taxonomy that had sampling of all cortical layers. The number of genes detected across tissue sources is highly correlated (**Fig. S3a**; R=0.99; p=3.3E-20), with a trend in neurosurgical cases toward higher gene detection in non-neuronal cells and lower gene detection in certain neuronal populations; however, variation in gene detection is approximately twice as high in neurosurgical cases. For neuronal types, most genes show consistent expression across tissue types. More variability is evident in non-neuronal types, where we find selective expression of a subset of genes in neurosurgical cases (**Fig. S3b**). Similarly, we find good agreement in the overall abundances of most cell types between tissue types (**Fig. S3c**). While we are underpowered to quantify differences in abundances, we do find relatively fewer Pvalb neurons in epilepsy cases than expected from the reference MTG data (**Fig. S3d**), which we follow up below by comparing epilepsy and tumor cases. However, variation in abundances is increased for nearly all cell types in neurosurgical cases compared to postmortem reference data (**Fig. S3e**). This increased variation might be due to differences in neurosurgical tissue dissections, as subclasses primarily located in the same layer tended to be strongly correlated (**Fig. S3f**). For example, L2/3 IT was correlated with superficial GABAergic types and L6 IT, L6b, and L6 CT types were strongly correlated. We also find anti-correlation between highly abundant types: astrocytes and oligodendrocytes were strongly anticorrelated and L2/3 IT was anticorrelated with other glutamatergic types. Overall, these results indicate consistent but more variable metrics in neurosurgical than postmortem tissue. While some of this variation can be explained technically, further analysis of population variation is needed.

### Population variation in cell type abundances

Next, we employed a linear model to investigate if demographic metadata, brain region, or donor medical conditions could explain the variance seen in gene counts and cell type abundances per cell type. Variability in gene counts was not significantly associated with any variable for any cell type (**Fig. S4**). In contrast, medical condition and brain region accounted for some abundance changes (**Fig. 2a–b**). For example, L5 ET neurons had lower abundance in tumor cases (FDR<0.01) even after accounting for significant regional differences in L5 ET cell types (FRO<MTG, FDR<0.01). In line with comparisons to the MTG reference taxonomy, Pvalb interneurons were slightly reduced in abundance in epilepsy vs. tumor cases (nominal p < 0.05). While this result is not significant after accounting for other covariates, the trend is consistent with previous reports demonstrating reduced Pvalb+ interneurons in certain focal cortical dysplasias (*37*) and decreased abundance of and gene expression changes in certain Pvalb+ interneurons (and other neuron types) in epilepsy (*38*). Additionally, L4 IT neurons were more abundant in MTG (FDR<0.01), while L6b neurons were more abundant in FRO (FDR<0.05). These differences were not associated with medical condition and likely reflect differences in cytoarchitecture between cortical areas. Although cell type abundances did not significantly differ by sex, some non-neuronal types showed a trend towards sex differences. For example, oligodendrocytes were slightly increased in males versus females (**Fig. S5a**; uncorrected p=0.18), consistent with previous reports showing an androgen-dependent increase in oligodendrocytes in male rats (*39*–*41*). Only OPCs show significant association with age, decreasing approximately two-fold across from ages 20 to 70 (**Fig. 2a; Fig. S5b**), consistent with a similar decrease in generation of OPC daughter cells in mouse hippocampus between 6 and 24 months (*42*).

Proper local and global brain network dynamics require a tight balance of excitation and inhibition (*43*), and loss of this balance can impact whole brain dynamics (*44*). Dysregulation of this balance is implicated in neurodevelopmental disorders, such as autism spectrum disorders, where an increased excitatory-inhibitory (E-I) ratio has been linked to memory, cognitive, and motor deficits and increased seizures (*45*–*47*). A breakdown of this E-I balance has been directly recorded in human during seizures (*48*), but whether such dynamics relate to differences in underlying cellular make-up isn’t currently known. To address this, we calculated the E-I ratio for each donor and applied the linear model described above (**Fig. S5c**). Consistent with previous work in primary motor cortex (*16*), the average E-I ratio in human was approximately 2, but the spread of E-I ratio is quite large across individuals (sd = 0.66), and this high variation was not associated with medical condition, sex, or brain region. These results do not confirm the hypothesis that E-I ratio is critical for network dynamics in this context.

### Cell type- and demographic-dependent donor signatures

Many sources of variation can impact gene expression within a subclass including higher resolution cell types (e.g., supertype), cortical cell depth (*23*), experience-dependent transcriptomic states (*34*, *49*), and disease state (*20*). To separate cell type from donor metrics, we calculated donor entropy, which measures the number of donors of origin for nuclei in a local neighborhood (**Fig. 2c**). Donor entropy is highly dependent on subclass, with nearly all GABAergic cells showing high donor entropy, and is also variable within subclass. For example, for most glutamatergic neurons and non-neurons, a subset of nuclei have high entropy while others have quite low entropy, consistent with visual inspection of **Fig. 1d**. Interestingly, OPCs and astrocytes with highly distinct transcriptomic signatures (low entropy island in **Fig. 2c**) were not technical artifacts and were dissected from two aged donors that had tumor resections and may represent reactive or pathological cells. To dissect the specifics of cell type and donor heterogeneity more carefully, we performed random forest (RF) classification (*50*) to predict donor of origin independently for nuclei in each subclass using the 2000 most highly variable genes. We find a wide range of RF prediction accuracies both by donor and by subclass (**Fig. 2d;** heatmap). Expression profiles of glutamatergic neurons are more predictive of donor overall than are profiles of most GABAergic and non-neuronal types (**Fig. 2d–e**). These results are robust to methodological details, as we find nearly the same results when predicting donor of origin using principal components (**Fig. S6a**). Similarly, RF predictability and the number of high variance genes show high concordance (**Fig. 2f**; p=7e-6), with both methods pointing to deep layer glutamatergic types as the most distinct types across this cohort.

RF prediction extends our gene variation analysis by allowing us to pinpoint specific donors with distinct gene signatures and allows us to ask whether these donors exhibit specific metadata characteristics. Interestingly, females show higher RF predictability than males in more than half of all subclasses including nearly all glutamatergic types (**Fig. 2d–e**). This can be partially explained by differential expression of sex chromosome genes, which have some of the highest importance for many donors in many subclasses (**Fig. S6b, 7**), in addition to being some of the most highly variable genes and having more RF importance in females than in males (**Fig. S6b-c**). Tissue from FRO and MTG tend to have comparable donor signatures, with the notable exception of several deep layer glutamatergic neurons (**Fig. 2d–e**). This may be due to selection bias, as tissue collected from MTG is relatively restricted anatomically, unlike tissue collected from spatially segregated regions of FRO, and glutamatergic neurons show larger gene expression differences across regions than GABAergic interneurons (*22*). In contrast to the abundance results, we do not see a difference in RF predictability by medical condition (**Fig. 2d–e**). This is consistent with previous reports demonstrating that, while tumor cells show large and tumor-specific changes in genomic copy number variations and associated gene expression signatures, nearby normal brain cells are well-mixed across samples (*51*, *52*). Overall, these results present a surprising sex difference and again point to layer 6 glutamatergic types as being highly distinct between individuals.

### Gene variation partitions into subclass and donor specific contributions

We have shown that neuronal and non-neuronal types exhibit unique patterns of variation in gene expression associated with donor (**Fig. 1d**, **Fig. 2c**). However multiple biological factors including ancestry (*53*), sex, age (*21*), and disease state (*20*) also contribute to the total variation in the expression of individual genes. To reveal the contribution of these biological as well as technical factors on transcriptomic variation we fit a subclass-specific linear mixed model per gene, termed variation partitioning (*54*). This analysis enables us to partition the per-gene total variance into the contributions from biological and technical factors as well as modeling residual variation that cannot be explained by the previous factors. Since the variance contributions per-gene sum to 1 it is straightforward to compare contributions across genes and subclasses.

We find individual genes and sets of genes whose variation is associated with biological factors in a subclass-specific manner (**Fig. 3a**). The number and magnitude of donor and supertype-associated genes ranges widely across subclass (**Fig. 3b**). Specifically, glutamatergic neurons have higher donor-associated contributions to variation than GABAergic and non-neuronal types which is in line with our quantification of donor entropy across neuronal classes (**Fig. 2c**). Within the glutamatergic types, we identified more donor-associated genes whose variation is explained more than 20% by donor within deeper layer neurons (n=1114) than L2/3 IT types (n=102), again pointing to these cell types as varying the most across individuals. The increased donor-associated genes in these deep layer types explains the improved performance of RF prediction on the task of predicting donor (**Fig. 2d**). Interestingly, GABAergic neurons show an increase in supertype-associated genes combined with lower donor-associated genes as compared to glutamatergic neurons (**Fig. 3b**). This is consistent with an increased supertype prevalence in GABAergic subclasses (**Fig. S8**) and coupled with limited GABAergic donor entropy implies an invariance with respect to donor in the expression of marker genes for GABAergic supertypes.

**Fig. 3.**
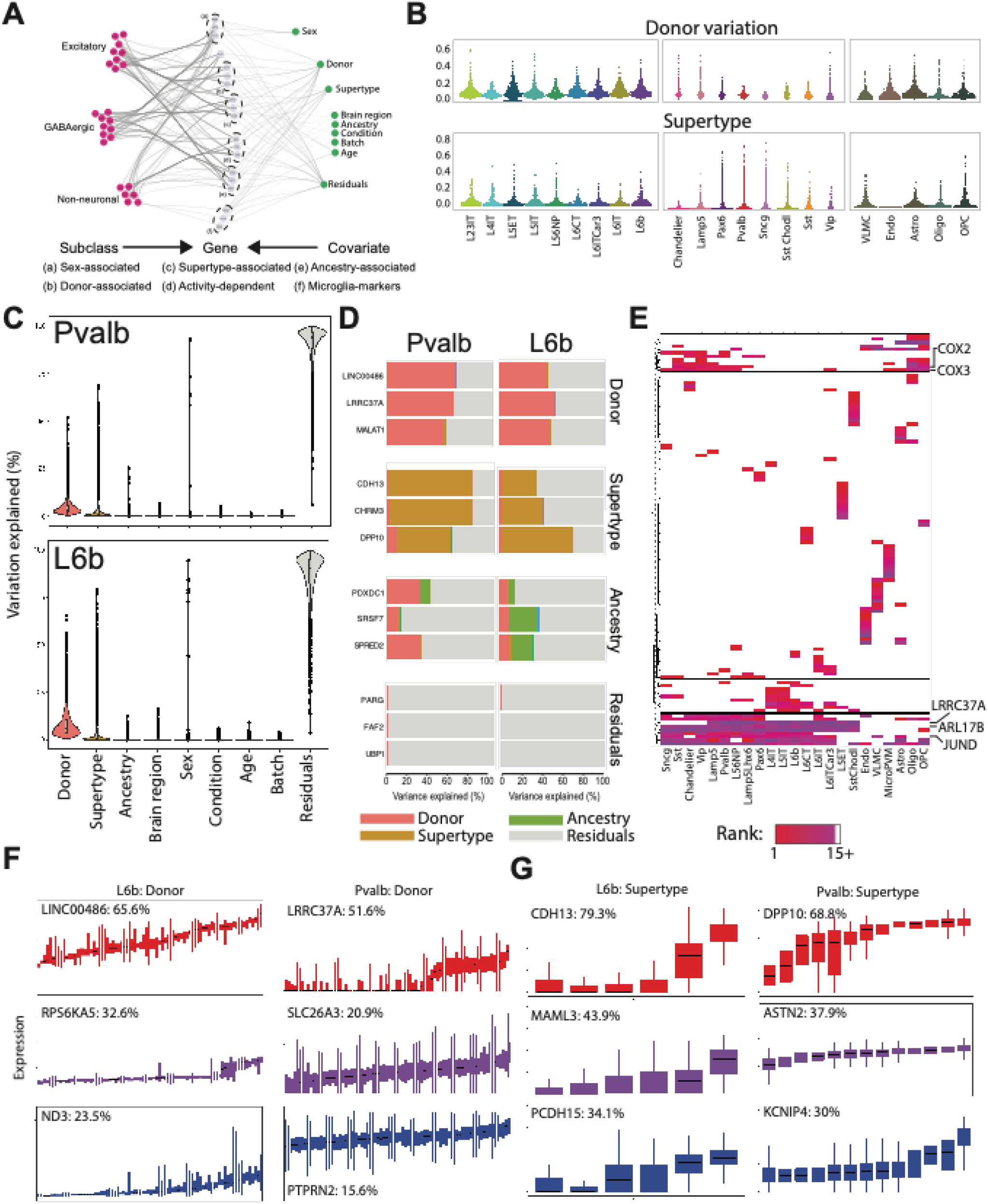
Variation partitioning explains differences in cell type-specific gene expression. (**A**) Interaction graph visualizing the estimated contribution of each covariate, purple node, to the variance in expression for each gene, green node, (covariate-gene edge) and the associated subclass, red node, (gene-subclass edge). The covariate-gene edge weight represents the precent variance explained for a gene by a covariate and the gene-subclass edge weight represents the non-residual contributions to variance explained for the gene. (**B**) Beeswarm plots showing for each subclass the amount of variation explained per-gene, represented by a point, by donor on the top row and supertype on the bottom row. (**C**) Violin plots showing the percentage of variation explained per-gene by each covariate for the Pvalb and L6b subclass. (**D**) Top genes whose variance can be associated with donor, supertype, ancestry and residuals for the Pvalb and L6b subclasses are shown in the paired boxplots. (**E**) Top ranked genes with respect to variance explained by donor across all subclasses are shown in the heatmap. Each row is a subclass, and each column is a gene which are colored based on rank determined from the percent variance explained by donor per-subclass. (**F**) Stratification of gene expression by donor for the top, median and 3^rd^ quantile donor-associated genes colored by red, purple and blue, respectively. Donors are ordered by median expression for each gene. (**G**) Stratification of gene expression by supertype for the top, median and 3^rd^ quantile donor-associated genes colored by red, purple and blue, respectively. Supertypes are ordered by median expression of each gene.

Transcriptomic variation across donors can be defined by known factors such as ancestry, sex and disease state and unknown factors that correspond to untracked or unidentifiable differences between donors. We know that some of the most highly variable genes across donors irrespective of subclass should be sex-associated genes including genes on the X and Y chromosome (**Table S2**). The variation partitioning analysis assigned nearly 90% of variation in sex-associated genes including *XIST*, *UTY* and *TTTY14* to the sex covariate as expected, especially *XIST* which is only expressed in females where it inactivates genes on one of the two X chromosomes (*32*) (**Fig. 3c**). Additionally, comparing the variation partitioning results across subclasses illustrates that variation between individuals is a major source of variation in gene expression, explaining >20% of variation in 52 genes expressed in Pvalb neurons and 336 genes expressed in L6b deep layer glutamatergic neurons (**Fig. 3c**). Of the top donor-associated genes we identified *LRRC37A* as being similarly highly variable in both Pvalb (51.6**%**) and L6b (63.1**%**) as well as genes including *HIPK1* and *GRB2* with donor-associated variability specifically in L6b neurons (**Fig. 3c–f**). With the exception of some pan-neuronal genes, the top donor-associated gene sets per subclass are largely distinct, likely due to the different gene programs which are active in each neuronal and non-neuronal types (**Fig. 3e**) (*17*, *55*). However, many of the most variable donor-associated genes are conserved across subclasses and associate with common gene ontology (GO) terms including cell-cell adhesion (FDR < 1e-15) and synaptic assembly (FDR < 1e-10).

Changes in gene expression variation with age have been associated with common pathways and disease development (*56*–*58*). Donors in this study varied in age from 18 years to 83 years and enable us to explore subclass-specific age-related gene programs. Age-associated changes in gene expression were identified from variance partitioning analysis to be largely subclass-specific (n=76/88, > 2% variation explained) and are predominantly contained in cell adhesion (FDR < 4.9e-4) and membrane (FDR <1.1e2) gene ontology categories. *ZBTB16* has been shown to vary significantly by age in studies of bulk-RNAseq in various tissues, including brain, from donors of comparable age ranges as this study(*57*). From our analysis we identified that age-associated variation in the expression of *ZBTB16* is oligodendrocyte-specific and refines previous findings to a specific cell type. Genes that exhibited age-associated variation across multiple subclasses included *PTPRD* which has been related to neurodegenerative disorders as well as roles in synapse maturation. Also, *LINC-PINT* had age-associated variation specific to excitatory subclasses and has been previously associated with various neurodegenerative diseases including Alzheimer’s disease (*59*).

Variation partitioning analysis is a useful tool for dissecting the contributions of biological covariates on gene expression. Yet, residual-associated genes remained the largest gene set in our analysis indicating that biological and technical covariates were not sufficient to completely explain gene variation patterns (**Fig.3c**, **Fig. S9**). Investigating genes whose variance is primarily explained by the model residuals, we found increased subclass-specificity. Such subclass-specific residual associated genes include gene markers such as *PVALB*, *SST*, and *SLC17A7*, which exhibit limited variation across donors within the associated types. Glutamatergic neuron residual-associated gene sets showed increased conservation across the more abundant types and include genes such as *ARID2* which is a known chromatin remodeling factor (*60*, *61*). Less abundant subclasses including L5 ET (n=563) and Sst Chodl (n=301) were enriched in subclass-specific residual-associated genes indicating gene programs uniquely active in these types with unknown sources of variation (**Fig. S9**). As a whole, these results indicate that transcriptomic variation associated with biological factors like donor and supertype can be cell type dependent, while also identifying sets of residual-associated genes whose variation cannot be explained by donor demographic information alone.

### Cell type-specific cis-eQTLs effects on gene expression variation

We next aimed to extend previous studies linking genetic variants to variation in gene expression (*24*–*27*, *62*–*64*) by identifying cell type-specific eQTLs from the combined single cell RNA-seq and high-quality whole genome sequencing (**Fig. S10**) performed for each individual in our study. We performed a cis-eQTL analysis for each subclass restricted to SNPs within a 1 megabase window surrounding gene transcription start sites (TSS) and adjusting for technical and inferred genotype covariates. Due to the limited number of donors, we restricted our analysis to SNPs associated with expression variation in previous bulk eQTL analysis (*24*, *25*, *60*). The number of significant cis-eQTLs (FDR < 0.05) varied widely across the subclasses with less abundant types including VLMC (n=637) and L6b (n=8,213) having relatively fewer cis-eQTLs than more abundant types such as L2/3 IT (n=79,631). Increases in cis-eQTLs for more abundant types indicates that sequencing more cells provides a better determination of variation within a subclass, up to a point. We find saturation in the number of detected eQTLs when at least 15,000 cells were sequenced in a given subclass (**Fig. 4a**). We identified an average of 200 eGenes, genes that are associated with a significant cis-eQTL, for each of the neuronal and non-neuronal subclasses (**Fig. 4b**), indicating that many cis-eQTLs were in linkage disequilibrium. Of the significant cis-eQTLs we noticed an enrichment of variants that are upstream of the TSS or around the gene body as compared to non-significant eQTLs (**Fig. 4c**). Of note, the non-neuronal type VLMC shows more cis-eQTLs downstream of the TSS which could imply differences in chromatin interaction patterns compared to the neuronal types.

**Fig. 4.**
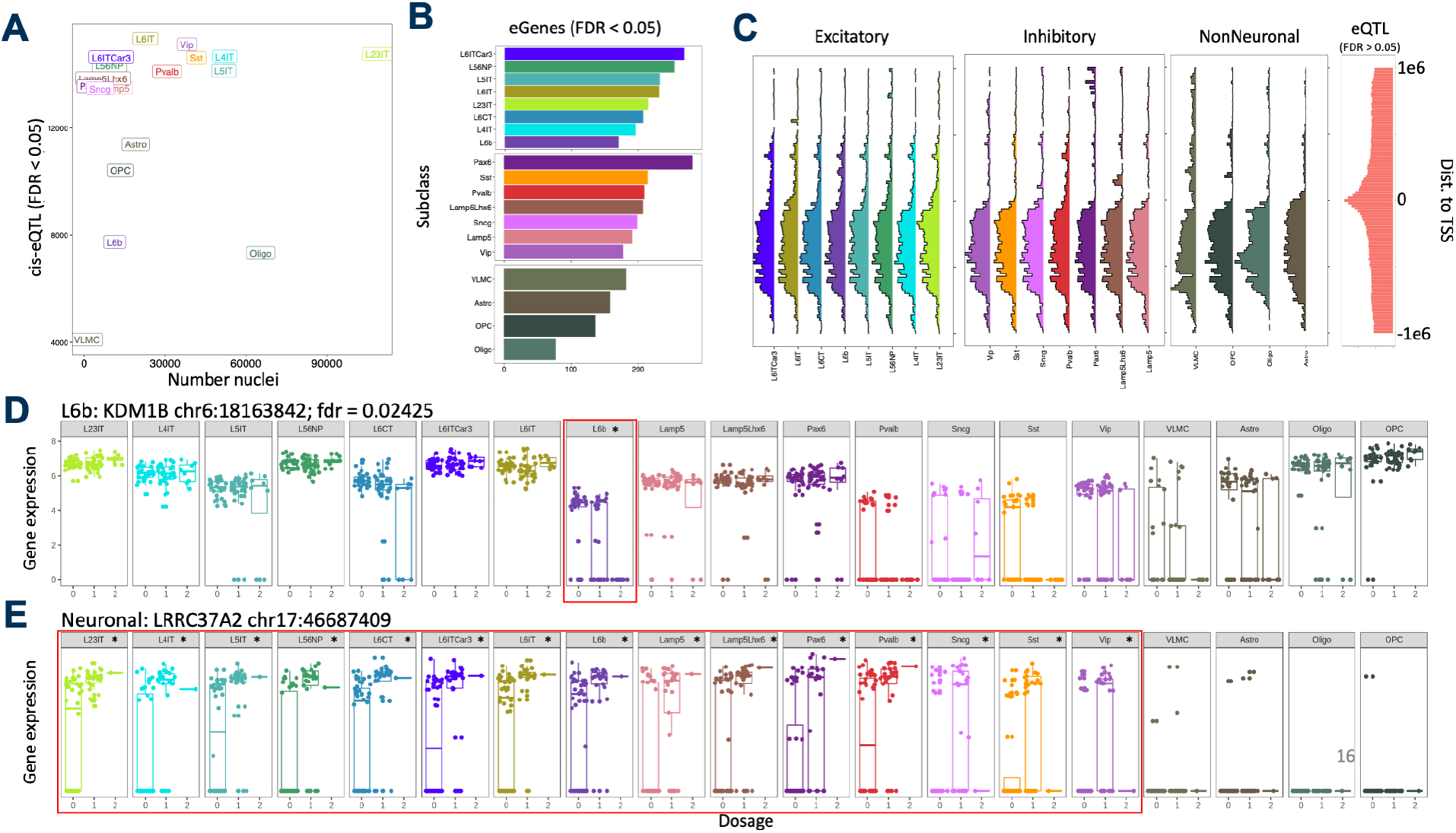
Cell-type specific gene expression variation is associated with genetic effects: (A) Scatterplot showing the relationship between numbers of cis-eQTLs (FDR < 0.05) on the y-axis and number of nuclei on the x-axis for each subclass. (B) Barplots show the amount of eGenes (FDR < 0.05) identified for each subclass, grouped by glutamatergic, GABAergic and non-neuronal. (**C**) Enrichment of cis-eQTLs upstream of the transcription start site of the proximal gene for each subclass compared to the background of all non-significant eQTLs (far right). (**D**) Example of a cis-eQTL, FDR < 0.05 denoted by *, that is specific to L6b neurons. (**E**) Examples of cis-eQTLs, FDR < 0.05 denoted by *, with effects on neuronal gene expression.

From the subclass-specific cis-eQTL analysis we next sought to understand the specificity of each cis-eQTL to gene expression variation for individual subclasses. Interestingly, we found cis-eQTLs which exhibited L6b and deep glutamatergic neuron specific control of gene expression variation including *KDM1B* (**Fig. 4d**). Additionally, genes including *LRRC37A2*, a paralog of and proximal to *LRRC37A* which we and others (*65*) previously identified as having donor-associated variation in expression, were identified as cis-eQTLs across neuronal cell types (**Fig. 4e**). This result indicates neuronal cell type-agnostic genetic control of LRRC37A2 which should recapitulate previous bulk RNA-seq eQTL studies. Together, these results show that genetic control of gene expression can extend to finer cell type resolutions than previously studied (*25*–*27*, *62*–*64*) and may be increasingly relevant for diseases including neurodegenerative disorders that may selectively affect specific cell types (*20*).

## Discussion

Here we present comprehensive transcriptomic and genomic profiling of cortical tissues from 75 human brain donors comprising nearly 400,000 nuclei covering all major neocortical cell types. Our results reveal that, while the cortical cellular architecture is largely conserved across individuals, substantial variation in gene expression and cell type abundances are apparent between individuals. Transcriptomic variation across donors is likely due to multiple competing factors. Underlying medical conditions in our donor cohort impact cellular abundance, with epilepsy cases showing decreased Pvalb interneurons, consistent with reports of fast-spiking interneurons loss in epilepsy (*37*, *38*). Variation in cellular gene expression is also apparent across neocortical areas, consistent with previous studies in human (*66*) and mouse (*22*). Similarly, L4 IT glutamatergic interneurons are more abundant in MTG than FRO, likely reflecting differences in cortical thickness and cytoarchitecture between regions (*67*). While we do not find sex-specific differences in cellular abundance, females do have more distinct gene expression profiles than males across many cell types, in part because these genes have a greater average RF score in females than in males (**Fig. S6c**) indicating their higher predictability in distinguishing female donors. Cellular pathways also dysregulate with age, leading to increased transcriptional heterogeneity in older vs. younger adults (*21*) and high variation in cell type abundances (*20*).

While we used largely nonpathological samples in the present study (*23*), tissue was nonetheless collected during neurosurgery; therefore, some variation is likely related to the circumstances of tissue collection rather than biological states. For example, immune response genes and markers for microglial states vary by donor, perhaps indicating differential responses to stress from surgeries (*68*) or underlying pathologies (*69*). Inconsistent tissue dissections could have major impacts on cell type abundances, as tissue shape or truncations might bias capture of cell types enriched in specific cortical layers, and the amount of white matter per section could impact abundance of oligodendrocytes. Such biases might explain the increased variability in this study relative to previous cell typing studies that made use of postmortem tissues, where the choice of dissection site is less constrained and more consistent between individuals (*17*). Future studies from postmortem cases with carefully controlled tissue sampling or *in situ* labeling of cell types using spatial transcriptomics methods could help to distinguish biological from technical variation.

Human glutamatergic neurons have higher inter-individual gene expression variability and predictability than GABAergic interneurons; but surprisingly, this variability is most prominent in deep layer glutamatergic types. Previous work showed that supragranular glutamatergic neurons are highly variable across many axes, including cell type, cortical depth, donor, and species (*16*, *17*, *23*), and UMAPs of these cells typically look less well mixed than for other glutamatergic types (**Fig. 1d**). Therefore, our initial expectation was that these L2/3 IT neurons would have the highest variation. However, deep layer glutamatergic neurons develop earlier (*70*) and have longer and more extensive connections (*71*) than superficial projection neurons. This exposes deep layer excitatory neurons and their projections to a more diverse environmental factors for a longer time during development than other neuronal types, potentially leading to more distinctive gene expression patterns across individuals with different life experiences, sensory inputs, and talents. Such donor variation provides opportunities to examine the association between gene expression and cellular function; for example, variability in ion channel gene expression might be linked with electrophysiological properties of cells from the same donors (*23*).

Some of the genes with highest inter-individual variability across subclasses relate to demographic information (e.g., sex chromosome genes like XIST, UTY, and TTTY14), brain region, or supertype (e.g., CDH13 in Pvalb interneurons). However, a much higher fraction of variance can be explained by other donor-associated factors. This includes activity-dependent genes (e.g., immediate early genes like JUND, FOS, and JUN) and genes marking nuclear content (e.g., MALAT1). Furthermore, the great majority of variation in all cell types is found in the residual component, reflecting unmeasured donor characteristics or technical variables (**Table S2**). Genes with such unexplainable variance should be avoided when targeting specific cell types *in situ* using spatial transcriptomic methods based on dozens to hundreds of selected genes (*72*). Instead, genes with high significance for subclass or supertype and low significance for donor metrics are likely to be robust to donor selection.

This study represents the most granular eQTL analysis to date where we identify genetic control of expression in subclass resolution cell types from adult human brains with coupled whole genome sequencing and single cell RNA-seq for 75 individuals. We report subclass-specific eQTLs providing an opportunity to associate disease-risk genes and variants from GWAS (*69*) with preferential regulation of expression in individuals or sets of subclasses. Current eQTL studies have used coarser groupings of cells than in this study and found eQTLs specific to excitatory and GABAergic neurons (*27*) that we can further resolved into subclass-level eQTLs to hone in more granular cell types targeted by variants associated with neurological disease. Furthermore, computational deconvolution (*14*, *15*) of eQTL analyses performed on bulk tissue (*26*, *62*, *73*) to resolve cell type signals can be refined using (*14*, *15*, *26*) the subclass-specific eQTLs reported in this study.

Two overlapping genes on chromosome 17, LRRC37A and ARL17B, showed high inter-individual variability selectively in neuronal but not glial types (**Fig. 3E**), and LRRC37A also contained many eQTLs for neuronal but not glial types (**Fig. 4E**). These genes are located at chromosome 17q21.31 alongside MAPT, which encodes tau protein, and are part of a common inversion polymorphism of approximately 900 kb that define the H2 haplotypes of MAPT (*65*). The H1 haplotype (no inversion) has been implicated in multiple neurodegenerative diseases through aggregation of hyperphosphorylated protein Tau in neuronal cell bodies (*74*, *75*). LRRC37A shows lower expression in temporal and frontal cortex of individuals with H1 vs. H2 haplotype in populations with European ancestry (*61*), suggesting a possible protective effect. In fact, a recent study on Parkinson’s disease linked protective sub-haplotypes of this locus with increased expression of LRRC37A in brain, although LRRC37A was primarily expressed in astrocytes in the substantia nigra (*76*). Although further work is needed to disentangle the effects of LRRC37A in neurons vs. astrocytes, and to see whether increased LRRC37A expression is likewise protective in other tauopathies, our results point to LRRC37A and ARL17B as important candidates for consideration in future studies of neurodegeneration.

This study has several limitations that represent opportunities for improvement in future population studies. First, the cohort primarily includes donors of European ancestry, representing the cross-section of the population undergoing neurosurgery in the local geographic region. Second, sampling neurosurgical tissues permits comparison of different underlying medical conditions between donors but introduces challenges in direct comparisons with similar studies using postmortem tissue and might result in biases due to tissue shape and sampling of variable cortical regions across donors. Third, the sample size is relatively small, although comparable results were found in a similar study in aging brain (*27*). Finally, the total number of cells collected per donor was relatively low (~4000 on average, collected from a single tissue section) limiting analyses possible for rare types.

In summary, this study provides a genomic and transcriptomic overview of cortical cell type variation in adult human individuals, identifying a highly consistent cellular makeup but with significant variation reflecting donor characteristics, disease condition, and genetic regulation.

## Acknowledgments

We thank the tissue procurement, tissue processing and facilities teams at the Allen Institute for Brain Science for assistance with the transport and processing of neurosurgical brain specimens and the technology team at the Allen Institute for assistance with data management. We thank our collaborators and hospital case coordinators at the Swedish Medical Center, Harborview Medical Center, and University of Washington Medical Center in Seattle for coordinating human neurosurgical tissue donations and tissue collections. We gratefully acknowledge the contributions of the donors who provided tissue samples that made this study possible. The content is solely the responsibility of the authors and does not necessarily represent the official views of the National Institutes of Health. The authors thank the founder of the Allen Institute, Paul G. Allen, for his vision, encouragement, and support.

## Funding

Allen Institute for Brain Science

National Institute of Mental Health of the National Institutes of Health grant U01MH114812 (ESL, TEB, RDH)

## Author contributions

Conceptualization: ESL, JAM, RDH, TEB

Investigation: AK, AMY, AT, BPL, CC, CDK, DH, DS, HT, JG, JG, JGO, JS, KS, MF, ND, NS, RDH, RE, RG, TC, TP

Formal analysis: JAM, KJT, MS, NJ, NLJ, SS, TEB Supervision: ESL, JAM, RDH, TEB

Writing – original draft: JAM, KJT, NJ, RDH, SS, TEB

## Competing interests

Authors declare that they have no competing interests.

## Data and materials availability

Raw snRNA-seq and WGS data is available from the NeMO archive after applying for controlled access through NDA and is listed under the project title: “A Multimodal atlas of human brain cell types: Human variation RNAseq & WGS (Lein)”. Code for reproducing the presented analyses will be made available prior to manuscript publication on GitHub (https://github.com/AllenInstitute/human_variation). Processed data will be made available for exploration and data download through CELLxGENE (*30*) and a link to these data will be included on GitHub prior to publication.

## Supplementary Materials

Materials and Methods

Figs. S1 to S10

Tables S1 to S4

References (*77*–*83*)

## Notes

### Competing Interest Statement

The authors have declared no competing interest.

