## Supplementary Materials for "Inter-individual variation in human cortical cell type abundance and expression"

**This PDF file includes:**

Materials and Methods
Figs. S1 to S10
Captions for Tables S1 to S4

**Other Supplementary Materials for this manuscript include the following:**

Tables S1 to S4 (zipped .xlsx file)

### Materials and Methods

#### *Experimental data collection*

Detailed descriptions of all experimental data collection methods in the form of technical white papers can also be found under 'Documentation' at http://celltypes.brain-map.org. (*23*)

*Human tissue acquisition and processing*

Tissue specimens were obtained from local hospitals (Harborview Medical Center, Swedish Medical Center, and University of Washington Medical Center) in collaboration with local neurosurgeons. A hospital-appointed case coordinator obtained informed consent from all donors (**Table S1**) before surgery and experimental procedures were approved by hospital institute review boards before commencing the study. The specimens collected for this study were apparently non-pathological tissues removed during the normal course of surgery to access underlying pathological tissues. Tissue specimens used in the study were determined by medical staff to be non-essential for diagnostic purposes and would have otherwise been discarded. Tissue specimens were de-identified before receipt by Allen Institute personnel. Immediately following resection, tissue was placed in artificial cerebral spinal fluid (ACSF) composed of (in mM): 92 N-methyl-d-glucamine chloride (NMDG-Cl), 2.5 KCl, 1.2 NaH2PO4, 30 NaHCO3, 20 4-(2-hydroxyethyl)-1-piperazineethanesulfonic acid (HEPES), 25 d-glucose, 2 thiourea, 5 sodium-l-ascorbate, 3 sodium pyruvate, 0.5 CaCl2.4H2O and 10 MgSO4.7H2O. Before use, the solution was equilibrated with 95% O2, 5% CO2 and the pH was adjusted to 7.3 by addition of 5N HCl solution. Osmolality was verified to be between 295–305 mOsm kg−1. Specimens were then transported (15–35 min) from the hospital site to the laboratory for further processing.

Acute brain slices (350 µm thickness) were prepared with a Compresstome VF-300 (Precisionary Instruments) or VT1200S (Leica Biosystems) vibrating microtome modified for block-face image acquisition (Mako G125B PoE camera with custom integrated software). Slices were transferred to oxygenated and warmed (34 °C) ACSF as described above for 10 min and were then transferred to room temperature holding ACSF composed of (in mM): 92 NaCl, 2.5 KCl, 1.2 NaH2PO4, 30 NaHCO3, 20 HEPES, 25 d-glucose, 2 thiourea, 5 sodium-l-ascorbate, 3 sodium pyruvate, 2 CaCl2.4H2O and 2 MgSO4.7H2O for at least two hours prior to freezing the slices for downstream nuclear isolation. Slices for RNA-sequencing were snap frozen in microcentrifuge tubes in a slurry of dry ice and ethanol and were stored at -80°C for later use.

*Nuclear isolation and single nucleus RNA sequencing*

Nucleus isolation for 10x Chromium Single Cell 3' RNA sequencing (v3) was conducted as described (*16*). Briefly, snap frozen tissue sections were removed from the -80°C freezer, thawed in homogenization buffer, and homogenized in a Dounce to generate dissociated nucleus suspensions. Nuclear suspensions were stained with DAPI to discriminate nuclei from debris and mouse anti-NeuN antibody conjugated to PE (Millipore, FCMAB317PE, 1:500 dilution) was applied to nuclear suspensions to discriminate neuronal (NeuN+) and non-neuronal (NeuN-) nuclei by fluorescence-activated nuclear sorting (FANS). Nuclei were sorted at a defined ratio of 80% NeuN+ neuronal and 20% NeuN- non-neuronal nuclei. After FANS, single-nucleus suspensions were concentrated by centifugation and frozen in a solution of 1X phosphate-buffered saline (PBS), 1% bovine serum albumin (BSA), 10% dimethylsulfoxide (DMSO) and

0.5% RNAsin Plus RNase inhibitor (Promega, N2611), and stored at −80 °C. At the time of use, frozen nuclei were thawed at 37 °C and processed for loading on the 10x Chromium instrument as described (*77*). Samples were loaded and processed using the 10x Chromium single-cell 3′ reagent kit v3 according to the manufacturer's protocol. For each donor, nuclei were loaded onto the 10x chip to target a maximum capture and sequencing of 10,000 single nuclei. Gene expression was quantified using the default 10× Cell Ranger v3 (Cell Ranger, RRID SCR_017344) pipeline, except for substituting of the curated genome annotation used for SMART-seq v4 quantification. Introns were annotated as 'mRNA', and intronic reads were included to quantify expression.

*10x Chromium Whole Genome Sequencing*

Tissue sections for whole genome sequencing (WGS) were from the same brain tissue block as used for snRNA-seq data generation. Immediately after slicing, tissue sections for WGS were mounted on glass slides, snap frozen, and stored at -80°C in 50ml conical tubes until later use. To isolate genomic DNA (gDNA), tissue was scraped off the glass slide using a sterile razor blade into a microcentrifuge tube and lysed overnight at 56°C using a Qiagen MagAttract HMW DNA Kit (Qiagen, 67563). Tissue lysates were processed according to the manufacturer's protocol, with several modifications as outlined in the 10x Genomics Genome Reagent Kit v2 User Guide. Final gDNA samples were run on an Advanced Analytical Fragment Analyzer using the Genomic DNA 50kb kit (Agilent, DNF-467) to measure gDNA concentration, assess the fraction of gDNA >40kb in each sample, and determine the quality of gDNA as measured using the genomic quality number (GQN). gDNA samples were diluted to 1ng/µl and concentrations were verified using the Qubit dsDNA HS Assay kit ahead of 10x loading. Samples were loaded and processed with the 10x Chromium Genome Chip Kit v2 and the Chromium Genome Library and Gel Bead Kit v2 using Chromium i7 Multiplex Kit sample indices. 10x Chromium sample processing followed the manufacturer's protocol. Samples were sequenced on a NovaSeq 6000 instrument using a NovaSeq S4 flow cell aiming for at least 25X genome coverage per sample.

*Variant calling in whole genome sequencing samples*

The processing of whole genome sequencing (WGS) data was done using SLURM on a high-performance cluster environment. The fastq files from 10x Chromium WGS were aligned using Burrows-Wheeler Aligner (BWA) mem to the human reference genome GRCh38-2.1.0. The bam files were processed using Picard (AddOrReplaceReadGroups) to add read groups. Variants were called on the processed bam files per chromosome using Genome Analysis Toolkit, GATK4-4.0.3.0-0 HaplotypeCaller with -ERC flag to get g.vcf files with homogenous reference calls. The individual g.vcf files were combined with GATK's CombineGVCFs and translated to vcf format with GATK's GenotypeGVCFs. Variants with phred-scaled quality score of 20 or greater were retained to ensure only high-quality variant calls.

***Reference MTG cell type definition and assignments for snRNA-seq data***

*Previously defined MTG reference cell type assignments*

All data, metadata, cell type assignments, associated taxonomy files, and documentation detailing protocols for each step are available at https://brain-map.org. In short, 151 clusters were defined as described in a recent study of conservation across great apes (*78*), using a combination of automated and manual QC, Leiden clustering using Seurat, and merging of clusters with insufficient evidence of differentially expressed genes. In addition to defining these high-resolution cell types, lower-resolution subclass and class assignments are defined as described previously for mammalian primary motor cortex and match the published interlex: (https://scicrunch.org/scicrunch/interlex) terms. Example subclass terms include "SST", "L6 CT", and "Astrocyte", while example class terms are one of "Neuronal: GABAergic", "Neuronal: Glutamatergic", or "Non-neuronal and Non-neural". All cells passing QC are assigned to the same class, and nearly all are assigned to the same subclass across taxonomies, even though independent cell type assignments are generated for each study.

*Creation of MTG reference supertype annotations*

We defined "supertypes" as a set of fine-grained cell type annotations for single nucleus expression data that could be reliably predicted on held-out reference data (where "ground truth" labels were assigned as described above) using state-of-the-art machine learning approach (*19*). From 5 neurotypical donors in the GA study with roughly 150K nuclei captured with 10x snRNAseq we systematically held out 2 donors at a time and used scANVI to iteratively and probabilistically predict their class (3 labels), subclass (24 labels), and then cluster (151 labels). When predicting each nucleus' class, we selected the top 2,000 highly variable genes along with the top 500 differentially expressed genes unique to each class (calculated from the reference cells which had their labels retained using a Wilcoxon rank sum test) to use as features in training the model and specified the donor name and number of genes detected as categorical and continuous covariates, respectively. Nuclei were then separated by their predicted class and features were re-selected with the same criteria to predict subclasses and again in predicting clusters. A differential expression test was run on clusters with an F1 score below 0.7, and those without 3 positive markers when compared against nuclei from their constituent subclass (corrected p-value <0.05, fraction in group expression >0.7, fraction out of group expression <0.3) were pruned from the taxonomy. Of the 26 clusters flagged, 24 fell below these cutoffs and were pruned from the final supertype taxonomy. The remaining 2 (L2/3 IT_2 and Oligo_3) were retained and recovered after supertype prediction (see below).

*Pre-mapping quality control of isolated nuclei*

Nuclei with fewer than 1000 genes detected and doublet score > 0.3 (*79*) were filtered upstream of supertype mapping. Additional quality control flags for each cell were determined using the public R package QCR (https://github.com/AllenInstitute/QCR_HVS).

*Mapping isolated nuclei to reference supertypes*

After defining supertypes in a neurotypical reference, we iteratively and probabilistically predicted class, subclass, and supertype for our nuclei using scANVI. Briefly, each nucleus's class was predicted after projection into a shared latent space with reference nuclei using models trained with 2000 highly variable genes and 500 differentially expressed genes per class. The highly variable and differentially expressed genes were derived from the reference data. scANVI was provided donor name and number of genes as categorical and continuous covariates, respectively. Nuclei were then split by predicted class, projected into a refined class-specific latent space where class-specific subclasses were predicted, and similarly for subclass-specific supertypes.

*Post-mapping quality control using QCR and scANVI*

The subclass-specific latent spaces were then used to compute two-dimensional uniform manifold approximation and projections (UMAPs) and the scANVI predictions were evaluated by known marker gene expression (using signature scores defined by differentially expressed genes in reference nuclei). In regions reference nuclei occupied there was strong agreement in signature gene expression with our nuclei, indicating accurate prediction. There was more variable expression in regions with poor reference support (which also had higher uncertainty in their predictions). These areas represented either droplets with ambient RNA, multiple nuclei, dying cells, or transcriptional states missing from the reference, unique to a donor or found only in disease. To triage these possibilities, we used the QCR R package (https://github.com/AllenInstitute/Quality_Control_for_scRNA-seq_data) to fracture the graph into tens to hundreds of clusters (called "meta-cells") using high resolution Leiden clustering and then merged the clusters based on differential gene expression. Clusters and meta-cells were then flagged and removed if they had low within-group doublet scores, or number of genes detected eliminating common technical sources of transcriptional heterogeneity.

*Expanding the reference taxonomy for non-neuronal cells*
With common technical axes of variation removed, we then sought to identify nuclei that were transcriptionally distinct from the reference and add them to our supertype taxonomy. We constructed a new latent space for each subclass using scVI (*80*), where the model was aware of the supertype prediction for each nucleus, gene dispersion was allowed to vary per supertype, donor name and sex were passed as categorical covariates, and the number of genes detected in each nucleus and the fraction of mitochondrial reads were passed as continuous covariates. Using the neighborhood graph from this latent space, we clustered the nuclei into tens to hundreds of groups and merged them based on differential gene expression using a python version of scrattch.hicat (https://github.com/AllenInstitute/scrattch.hicat). We defined merged clusters with fewer than 10% of all reference cells or of any single supertype as having limited reference support and added them to the taxonomy named as: Subclass_Unknown_ClusterNumber.

***RNA-seq data analysis***

*Assessing gene variability across donors*
Differentially expressed genes between cell types or donor demographics (e.g., epilepsy vs. tumor) are calculated on log2 normalized counts per million using the Wilcoxon test using the FindMarkers function in the Seurat R library (*81*). Genes with higher within-donor than between-donor variance were identified as follows: 1) for each subclass, select (up to) 24 cells per donor; 2) calculate that average gene expression variance within each donor; 3) repeat the first two steps 11 times for different sets of cells with real and permuted donor IDs; and 4) compare the median real vs. permuted variances. Genes with the highest ratio of permuted to real variation have the largest donor effects and represent genes likely to be identified using variance partitioning (see below). Resulting differential expression and high variance gene sets were tested for enrichment of gene ontology (GO) and other categories using the ToppGene Suite (*82*).

*Assessing cell type variability across donors*
Cell type abundances were calculated separately for neuronal and non-neuronal cell populations to account for biases intentionally introduced through FANS, with abundances in each group summing to 100 percent per donor. Changes in abundances between donor demographics (e.g.,

epilepsy vs. tumor) were calculated using an ANOVA corrected for multiple comparisons using the false discovery rate method.

*Random forest prediction of donor ID*

Random forest prediction was used to assess the amount of intra-donor vs. inter-donor variation within each subclass by asking for what fraction of cells can the assigned cell type be accurately predicted. The specific algorithm is as follows: 1) for each subclass, select (up to) 24 cells per donor; 2) run random forest prediction (*50*) using 75% training, 25% test four times to get predictions for all 24 cells; and 3) repeat first two steps 10 times to calculate mean and standard deviation of RF prediction scores per subclass, per donor. These classifications accuracies are also calculated in data where donor IDs are randomized within subclass for comparison. Donors and subclasses are then hierarchically clustered by average prediction accuracy, groupings of subclasses and donors with respect to metadata are assessed using a multivariate hypergeometric distribution.

*Assessing donor entropy across cell types*

Donor entropy per subclass was used to assess the amount of alignment between donors. The nearest 100 neighbors for each cell was computed using `spatial.cKDTree` from the scipy library. The donor annotation for each of the 100 nearest neighbors were summarized as proportions that were used to compute donor entropy with the `stats.entropy` function from the scipy library. We repeated this procedure for each cell in each subclass.

*Variance Partition analysis*

Variation partitioning analysis was utilized to prioritize the drivers of variation within each subclass. Using linear mixed-effect models implemented in the variancePartitioning bioconductor package: http://bioconductor.org/packages/variancePartition (*54*) we identify the genes whose variance is best explained by donor, supertype, brain region, demographic (ancestry, age, sex) factors, disease states (epilepsy vs. tumor) and technical factors (batch). The specific algorithm is as follows: (1) for each subclass we filter to cells passing QCR flags and remove any clusters with a single member. (2) Genes are removed from the analysis if the gene was not expressed in > 10 cells, have greater than 80% dropout, have zero variance, have an expression of less than 1 CPM on average and are in the ZNF or LOC families. (3) The variance partitioning linear model was then defined as:

$$gene \sim age + (1|condition) + (1|sex) + (1|donor) + (1|supertype) + (1|batch) + \left(1\middle|brain_{region}\right) + (1|ancestry)$$

and passed into the variancePartition function `fitVarPartModel`. We determined the amount of variation explained per covariate for each gene with the `extractVarPart` function.

***Whole-genome sequencing analysis***

*eQTL analysis*

For eQTL analysis we filtered to the following SNPs: (1) minor allele frequency (MAF) < 5%, (2) significant in an eQTL from ROSMAP, CommonMind or brain tissue from GTEx. The .vcf files containing SNP data were converted to dosage files using samtools -dosage plugin. The gene expression matrix was filtered as follows: (1) not expressed in > 10 cells, (2) having greater than 80% dropout, (3) having zero variance, (4) having an expression of less than 1 CPM on average and (5) in the ZNF or LOC families. After filtering both the SNP and gene expression data we used the Matrix eQTL software package (*82*) to identify significant eQTLs. In the Matrix eQTL model we included the top 3 genotype PCs to account for ancestry differences as well as technical covariates including the tissue collection site.

*Acquiring ROSMAP eQTL summary statistics*
We obtained eQTL summary statistics for the ROSMAP study from the xQTL analysis performed in Ng et al. 2017. We downloaded the summary tables for significant eQTLs from the updated xQTL statistics (2021) hosted on http://mostafavilab.stat.ubc.ca/xqtl/.

*Acquiring CommonMind eQTL summary statistics*
We obtained eQTL summary statistics for the CommonMind study from the CommonMind Consortium Knowledge portal Synapse project:
https://www.synapse.org/#!Synapse:syn4622659.

*Acquiring GTEx eQTL summary statistics*
We obtained eQTL summary statistics for brain tissues from version 8 of the GTEx study at:
https://www.gtexportal.org/home/datasets where we downloaded the following file:
https://storage.googleapis.com/gtex_analysis_v8/single_tissue_qtl_data/GTEx_Analysis_v8_eQTL.tar.

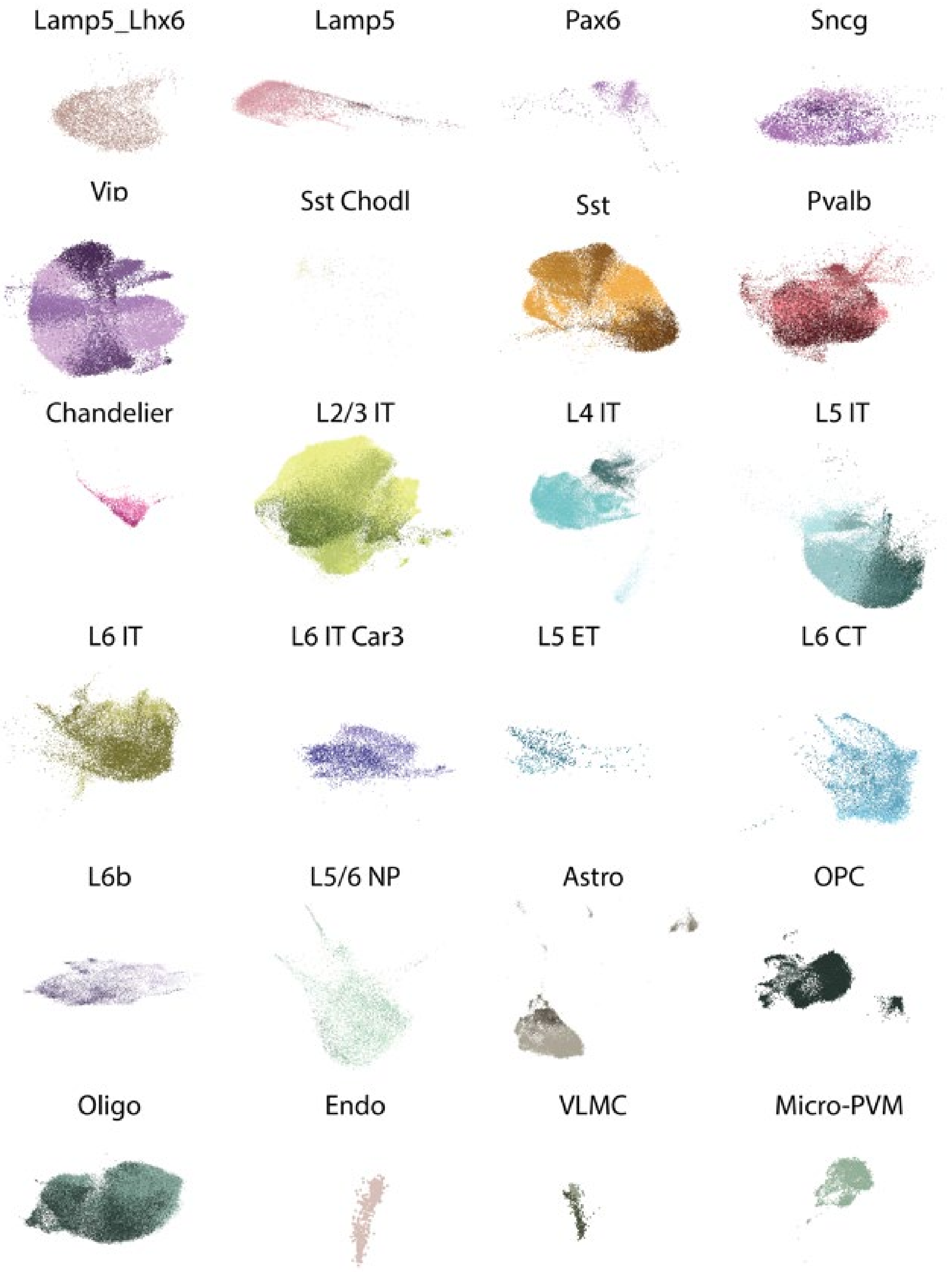


**Fig. S1.**
**UMAP of supertypes**. UMAP plot per subclass where each cell is colored by supertype annotation.

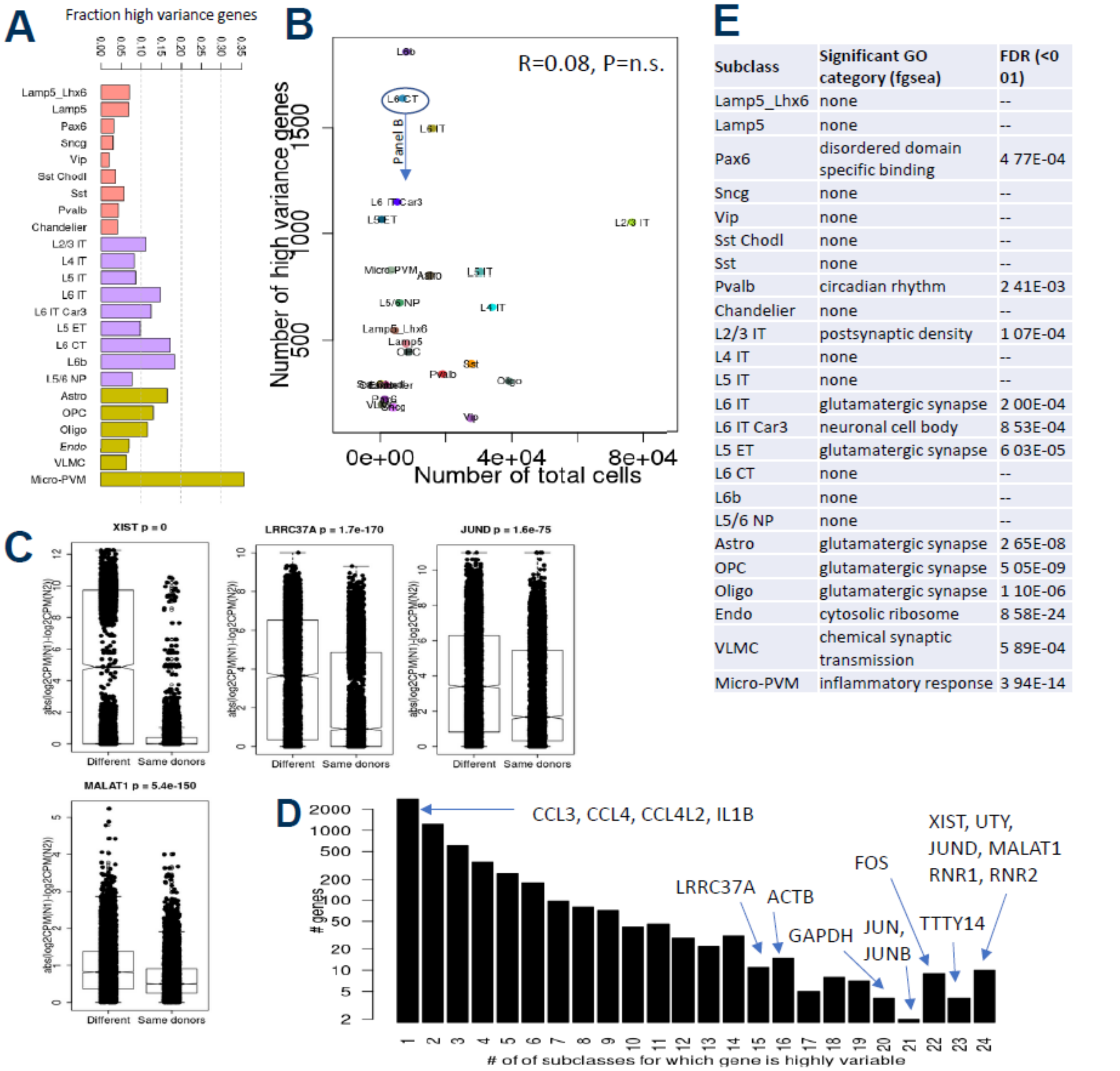


| Subclass | Significant GO category (fgsea) | FDR (<0 01) |
|---|---|---|
| Lamp5_Lhx6 | none | -- |
| Lamp5 | none | -- |
| Pax6 | disordered domain specific binding | 4 77E-04 |
| Sncg | none | -- |
| Vip | none | -- |
| Sst Chodl | none | -- |
| Sst | none | -- |
| Pvalb | circadian rhythm | 2 41E-03 |
| Chandelier | none | -- |
| L2/3 IT | postsynaptic density | 1 07E-04 |
| L4 IT | none | -- |
| L5 IT | none | -- |
| L6 IT | glutamatergic synapse | 2 00E-04 |
| L6 IT Car3 | neuronal cell body | 8 53E-04 |
| L5 ET | glutamatergic synapse | 6 03E-05 |
| L6 CT | none | -- |
| L6b | none | -- |
| L5/6 NP | none | -- |
| Astro | glutamatergic synapse | 2 65E-08 |
| OPC | glutamatergic synapse | 5 05E-09 |
| Oligo | glutamatergic synapse | 1 10E-06 |
| Endo | cytosolic ribosome | 8 58E-24 |
| VLMC | chemical synaptic transmission | 5 89E-04 |
| Micro-PVM | inflammatory response | 3 94E-14 |

**Fig. S2.**

**High-variance genes include known sex and activity-dependent genes but are otherwise largely diverse.** **A)** Fraction of high-variance highlights microglia. Plot is the same as Fig 1e (left), but scaled to the number of expressed genes per subclass. **B**) No relationship between the number of high variance genes (y-axis) and the number of cells per subclass (x-axis), suggesting the metric for determining high-variance genes is reasonable. **C**) Four example genes showing higher inter- vs. intra-donor variability in L6 CT. Bar plots showing the difference in log2CPM from 1000 randomly selected cells (x-axis) from the different vs. the same donors (x-axis). These genes are all highly variable in 15 or more subclasses (not shown). **D**) Inverse relationship between number of significant subclasses a gene is highly variable in (x-axis) and the number of genes in that group (y-axis). Some genes of interest are shown, including microglial markers (CCL3, CCL4, CCL4L2, IL1B) and the more ubiquitous genes of interest (sex genes, immediate early genes, etc.). **E**) One of the most significantly-enriched gene ontology categories per subclass. Several neuronal cell types did not have any significant categories.

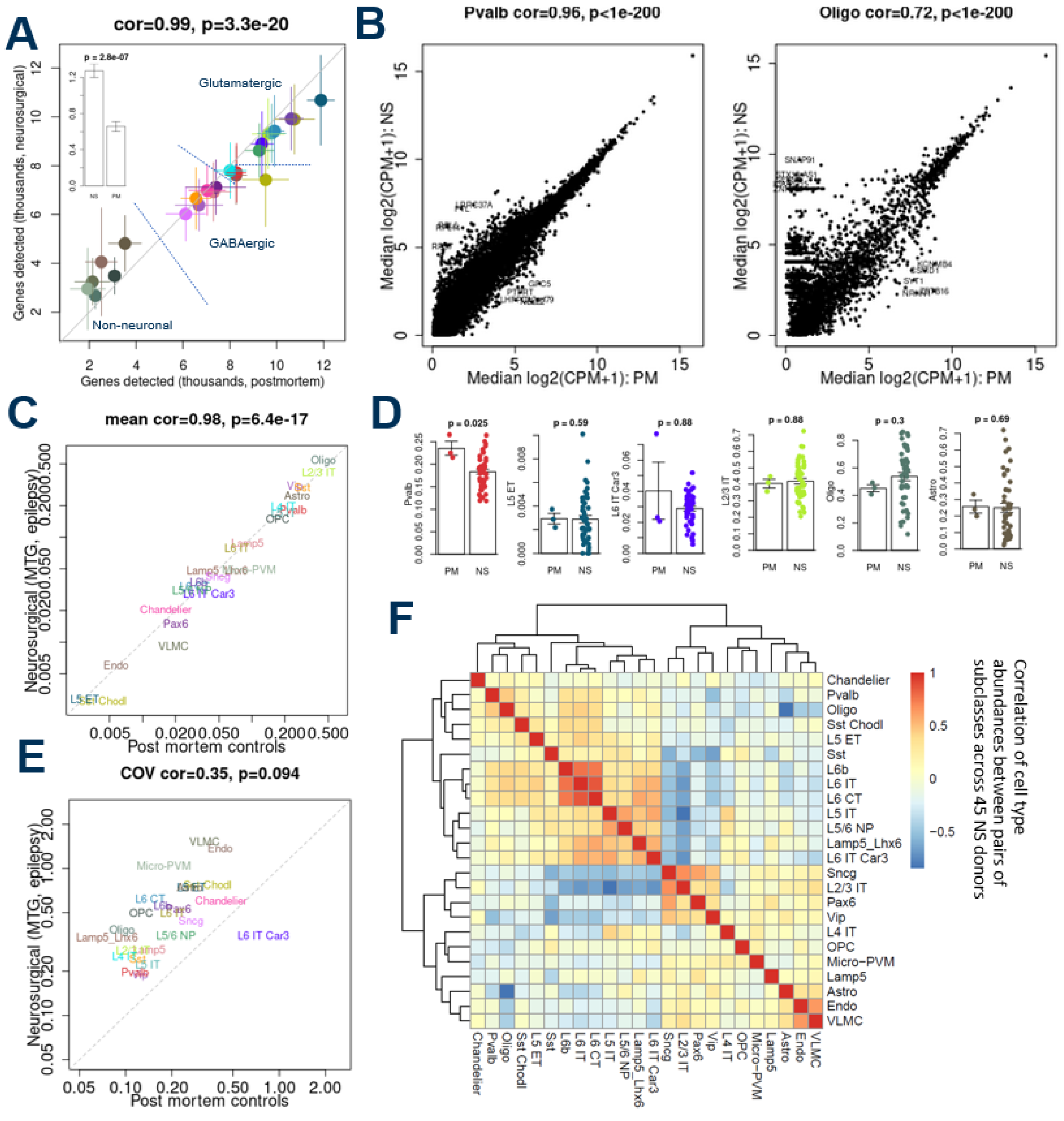


**Fig. S3.**

**Cell types and gene expression largely but not entirely conserved between neurosurgical and postmortem tissues A)** Number of genes detected per donor is high correlated between the 45 neurosurgical (NS) cases from MTG of epilepsy donors (y-axis) and the three postmortem (PM) reference donors with unbiased laminar sampling (x-axis) Inset: Significantly higher variation in the number of genes detected per donor (y-axis) in NS vs PM tissues **B)** High correlation in gene expression levels from the same NS vs PM cases, with neuronal cell types (e g , Pvalb, left) showing generally better agreement than non-neuronal cell types (e g , Oligodendrocyte, right) **C)** Good agreement in abundance levels between NS and PM tissues **D)**

(left) Significantly fewer Pvalb cells in NS than PM tissues, likely reflecting loss of Pvalb-interneurons epilepsy (Other plots) No difference in abundance but higher variation in NS vs PM tissues for other cell types, except for L6 IT Car3, which likely has higher variation in PM due to a single outlier donor **E**) NS donors have higher cell type variation than PM for all types except L6 IT Car3 **F**) Correlation of subclass abundances across NS cases match expectations (e g , L2/3 anticorrelated with other glutamatergic types but correlated with superficial inhibitory types, astrocytes and oligos strongly anticorrelated, and L6 types are strongly correlated, potentially reflecting dissection differences).

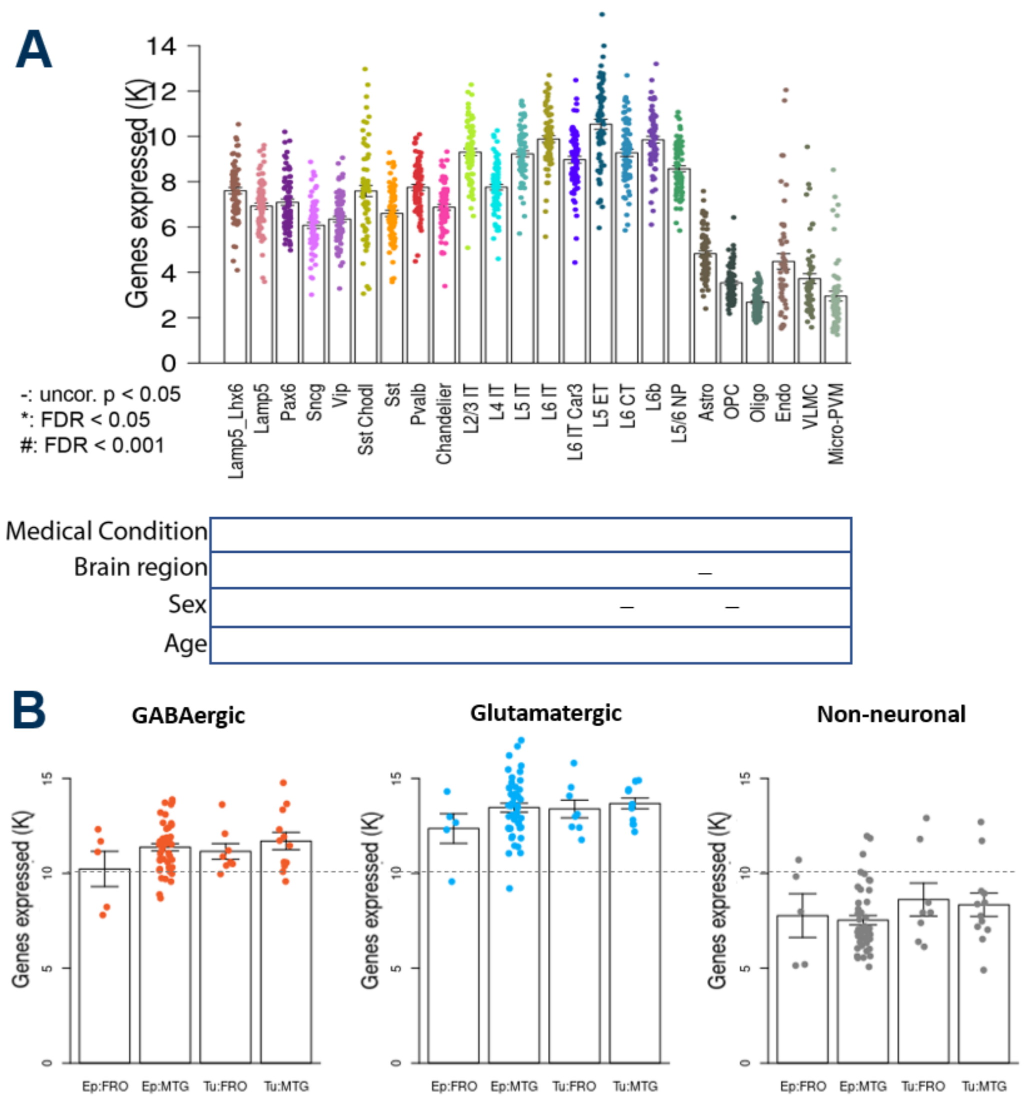


**Fig. S4.**

**Gene counts consistent with reference and unrelated to metadata**   **A)** Number of genes expressed per subclass is relatively consistent across donors and not significantly related to any metadata assessed after accounting for multiple comparisons   Labeling as in **Figure 1A**  As expected, more genes are expressed in excitatory than inhibitory neurons and more in neurons than non-neurons   Donors with exceptionally high numbers of genes expressed in non-neural types likely represent doublets or other imperfect QCing of the data  **B)** Number of genes expressed per donor per class   No significant differences across region or medical condition (p=NS for all three plots).

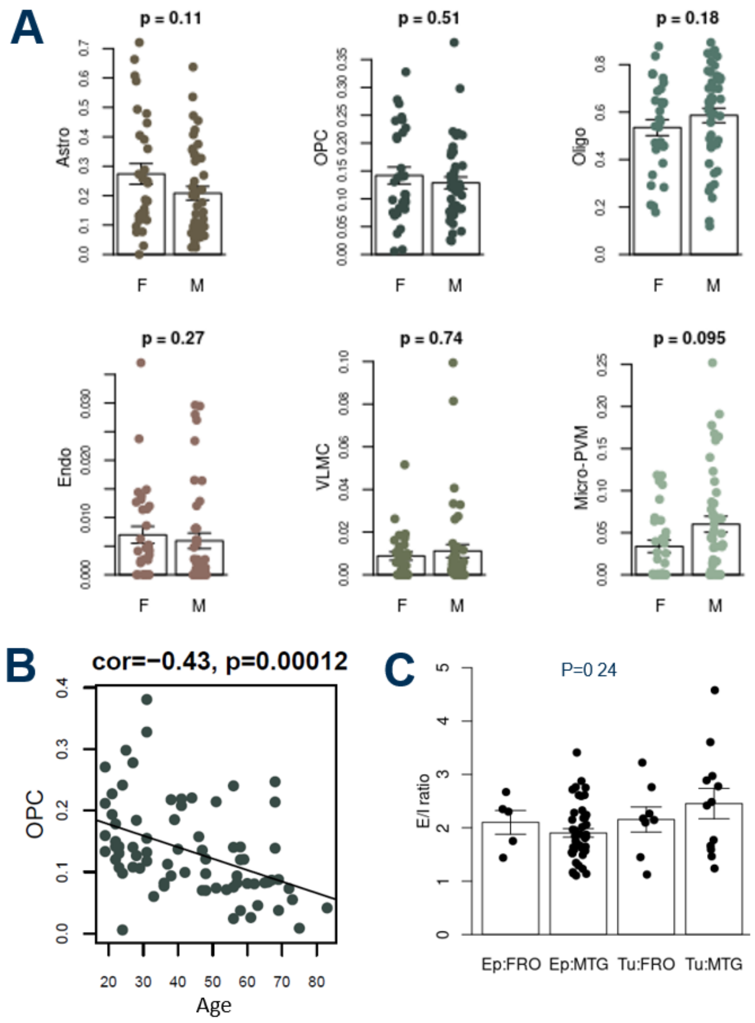


**Fig. S5.**

**Changes in cell type abundance by sex, age, region, and condition  A)** Non-neuronal cell types show some trends in abundances (y-axis) between males and females (x-axis), but no significant differences after accounting for covariates and multiple comparisons  P values indicate uncorrected Kruskall-Wallace test p values. **B)** OPCs show significant decrease in abundance (y-axis) with respect to age (x-axis)   P values indicate uncorrected correlation-based p values. **C**) Ratio of the abundance of excitatory to inhibitory cells per donor (E/I ratio; y-axis), separated by brain region and medical condition (as in **Figure 2b**)   Average ratio matches expectations, and not differences across metadata, but variation is high than  in previous studies.

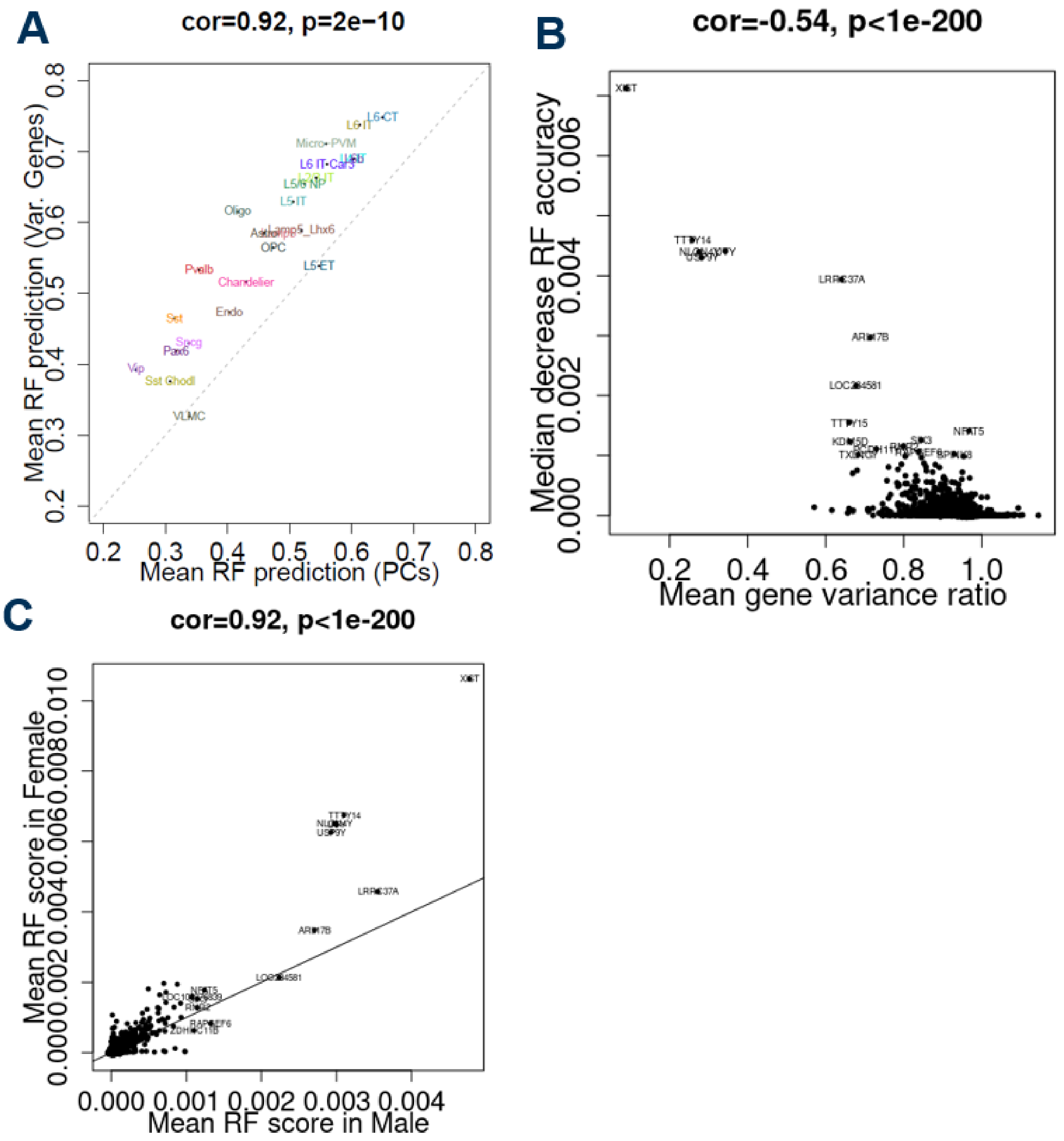


**Fig. S6.**

**Additional insights from RF prediction analysis A)** High correlation between median RF prediction accuracies defined from the top 2000 variable genes (y-axis; **Fig 2d**) and those defined using the top 30 principal components (x-axis) Prediction accuracies generally higher based on variable genes **B**) Highly variable genes (low values in x-axis, ratio of values from **Fig 2e**) also tend to be the ones most important in determining RF accuracy (high values in y-axis, which represents the mean decrease in accuracy per donor when removing that gene from the RF analysis) **C**) High correlation between RF prediction accuracy in male (x-axis) and female (y-axis), but sex genes are generally more predictive in female This may be due to their being fewer females than males in the study.

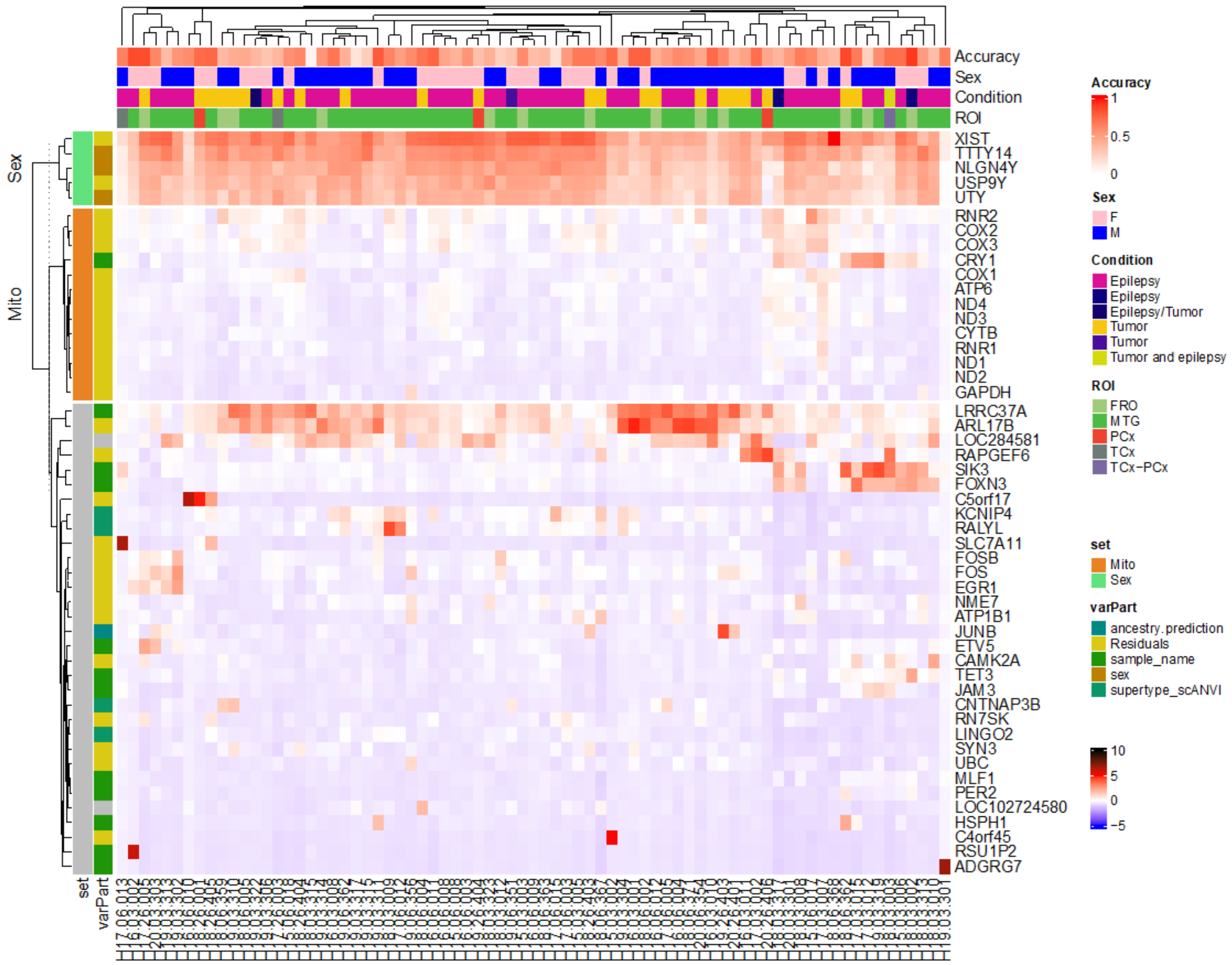


**Fig. S7.**

**Genes with the highest RF importance include sex, mitochondrial, and other genes**
Heatmap showing the genes with the highest RF importance in Pvalb subclass, with the specific accuracy shown per donor (x-axis) per gene (y-axis)   For each gene the category of highest partitioned variance (from **Figure 3**) is shown  With all sex genes showing high variance explained by sex or donor   Donors do not cluster by any of the metadata (medical condition, region, and sex) shown on the y-axis   Similar results (including many of the same genes) are identified across other subclasses (not shown).

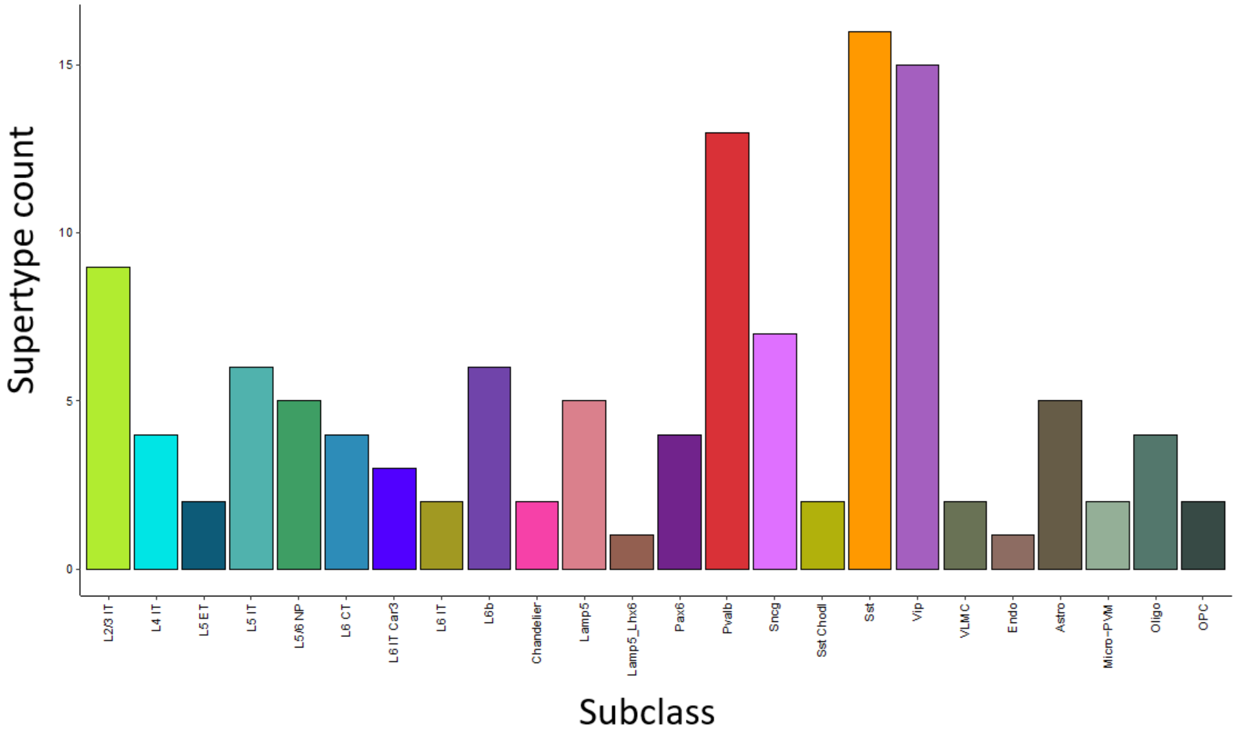


**Fig. S8.**

**Abundance of supertypes for each subclass.** Bar plot visualizing the number of supertypes (y-axis) per subclass (x-axis).

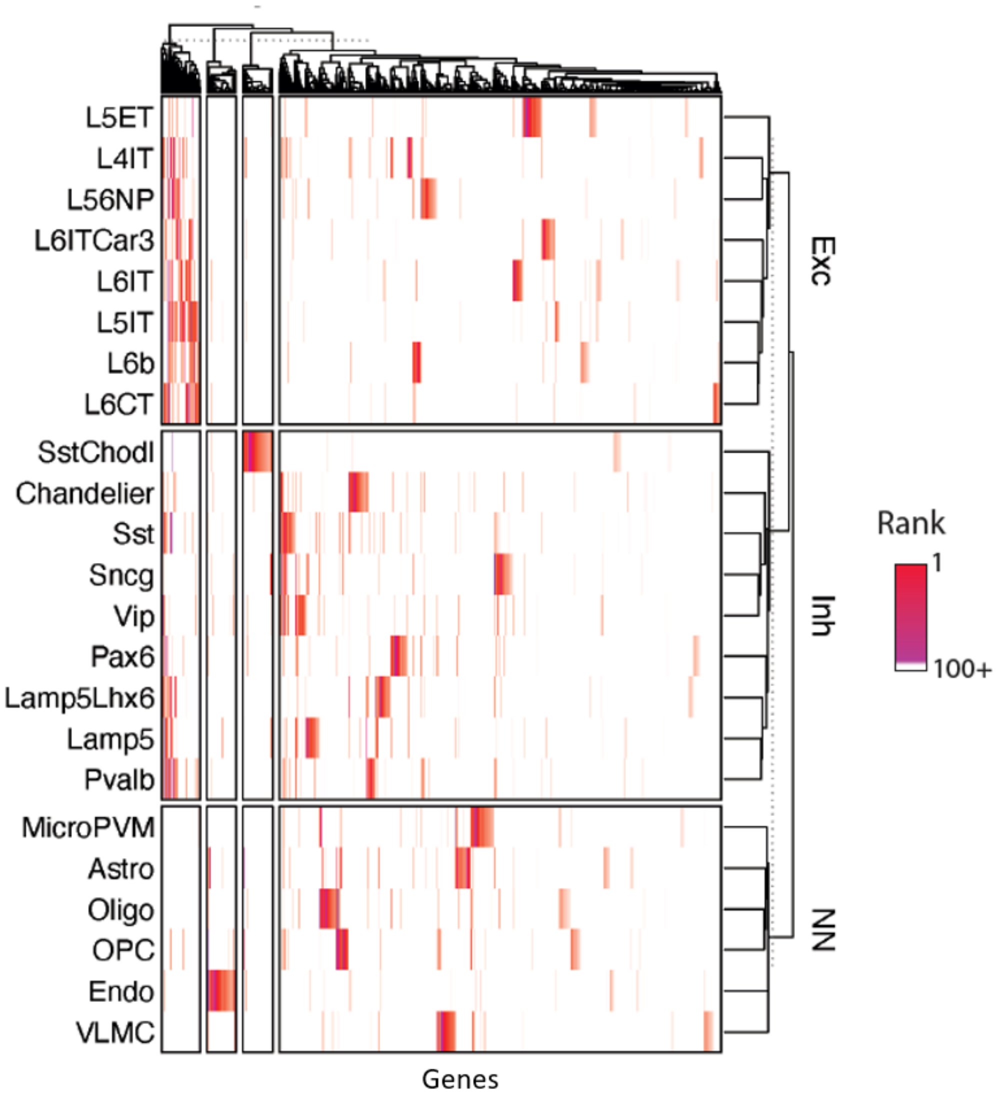


**Fig. S9.**

**Top residual-associated genes per subclass** Heatmap showing the top 100 residual-associated genes, per subclass, as defined from the variance partitioning analysis. Subclasses are grouped by neuronal class on the y-axis and genes are shown on the x-axis.

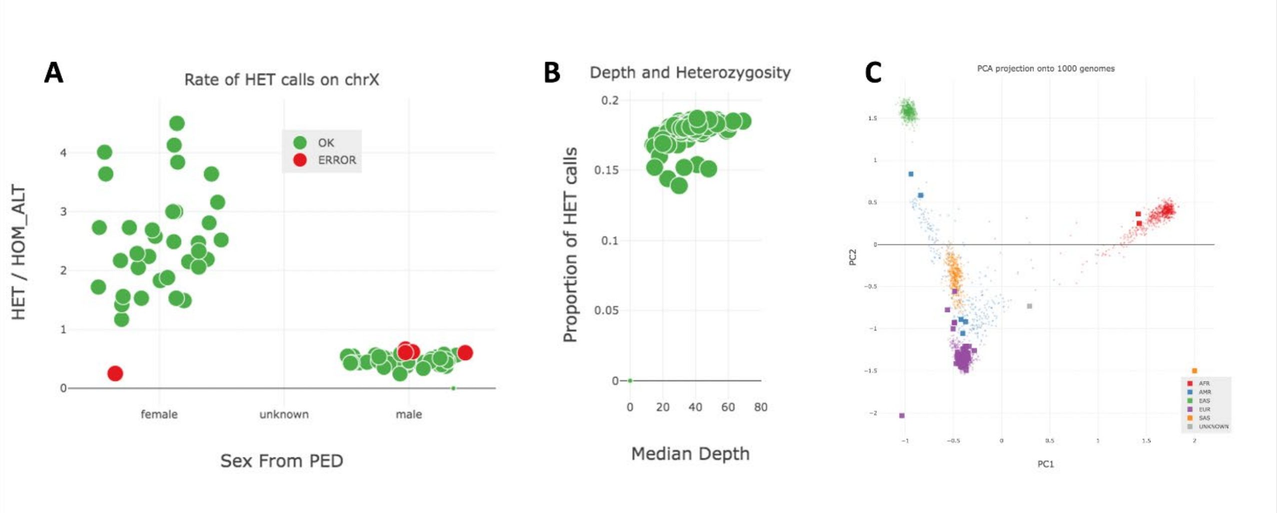


**Fig. S10.**

**Quality control and ancestry prediction using Peddy A)** Scatter plot visualizing on the x-axis the **s**ex from pedigree (PED) and on the y-axis the ratio of heterozygous (HET) to homozygous (HOM) genotypes from variants outside of pseudo-autosomal region of the X chromosome. Errors due to low coverage are indicated by red circles. **B)** Depth vs heterozygosity plot from Peddy to detect problems with DNA quality and purity and unexpectedly high levels of homozygosity/heterozygosity. Samples with green dots are OK, red dots are ratio outlier and blue dots are depth outlier. **C)** PCA showing the predicted ancestry of the samples using SVM trained on the 1000 genomes samples indicated by smaller dots in the background.

### Supplemental Tables

*Supplemental tables are all located in a separate zipped .xlsx file.*

**Table S1.**
Study metadata including donor demographic information and ancestry predictions.

**Table S2.**
List of sex-associated genes used in this study.

**Table S3.**
Complete variance partitioning results for all subclasses and genes analyzed.

**Table S4.**
MAPT haplotypes and LRRC37A, ARL17B and KDM1B genotypes for each donor.
